# Computational and biochemical analysis of type IV Pilus dynamics and stability

**DOI:** 10.1101/2021.04.14.439816

**Authors:** Yasaman Karami, Aracelys López-Castilla, Andrea Ori, Jenny-Lee Thomassin, Benjamin Bardiaux, Therese Malliavin, Nadia Izadi-Pruneyre, Olivera Francetic, Michael Nilges

## Abstract

Type IV pili (T4P) are distinctive dynamic filaments at the surface of many bacteria that can rapidly extend, retract and withstand strong forces. T4P are important virulence factors in many human pathogens, including Enterohemorrhagic *Escherichia coli* (EHEC). The structure of the EHEC T4P has been determined by integrating Nuclear Magnetic Resonance (NMR) and cryo-electron microscopy data. To better understand pilus assembly, stability and function, we performed a total of 108 μs all-atom molecular dynamics simulations of wild-type and mutant T4P. Extensive characterization of the conformational landscape of T4P in different conditions of temperature, pH and ionic strength was complemented by targeted mutagenesis and biochemical analyses. Our simulations and NMR experiments revealed a conserved set of residues defining a novel calcium-binding site at the interface between three pilin subunits. Calcium binding enhanced T4P stability *ex vivo* and *in vitro*, supporting the role of this binding site as a potential pocket for drug design.

## INTRODUCTION

In bacteria, multiple molecular machineries perform a wide variety of biological functions necessary for their survival, social organization, motility, and capacity of infecting host cells. Bacterial interactions with their environment often rely on surface polymers called pili, which are assembled through a regular repetition of one or few protein subunits called pilins. In some cases, pilins are irreversibly or covalently linked to each other, building fibers with high stability and mechanical resistance. This is the case of type I pili from Gram-negative (Hospenthal, Costa, and Waksman 2017) and sortase-dependent pili from Gram-positive bacteria (Kang and Baker 2012). Other pili are maintained by noncovalent interactions between subunits to combine mechanical strength with flexibility and dynamics. Remarkable representatives of this class are type IV pili (T4P), most notably the T4aP subclass that are able to rapidly assemble and disassemble (Craig, Forest, and Maier 2019). T4P dynamics is crucial for their biological functions, which include adhesion to host cells, twitching motility, DNA uptake, protein secretion, and microcolony formation. These fibers can reach several micrometers in length and range between 6-9 nm in diameter. The machinery that assembles T4P is formed by several protein sub-complexes that span the bacterial envelope (**Figure 1a**). T4P are anchored in the inner membrane (IM) and extend beyond the outer membrane (OM) of Gram-negative bacteria through a channel called the secretin. Dedicated ATPases at the cytoplasmic base of the complex transmit motions to the assembly platform IM complex to drive fiber extension and retraction.

**Figure 1.**
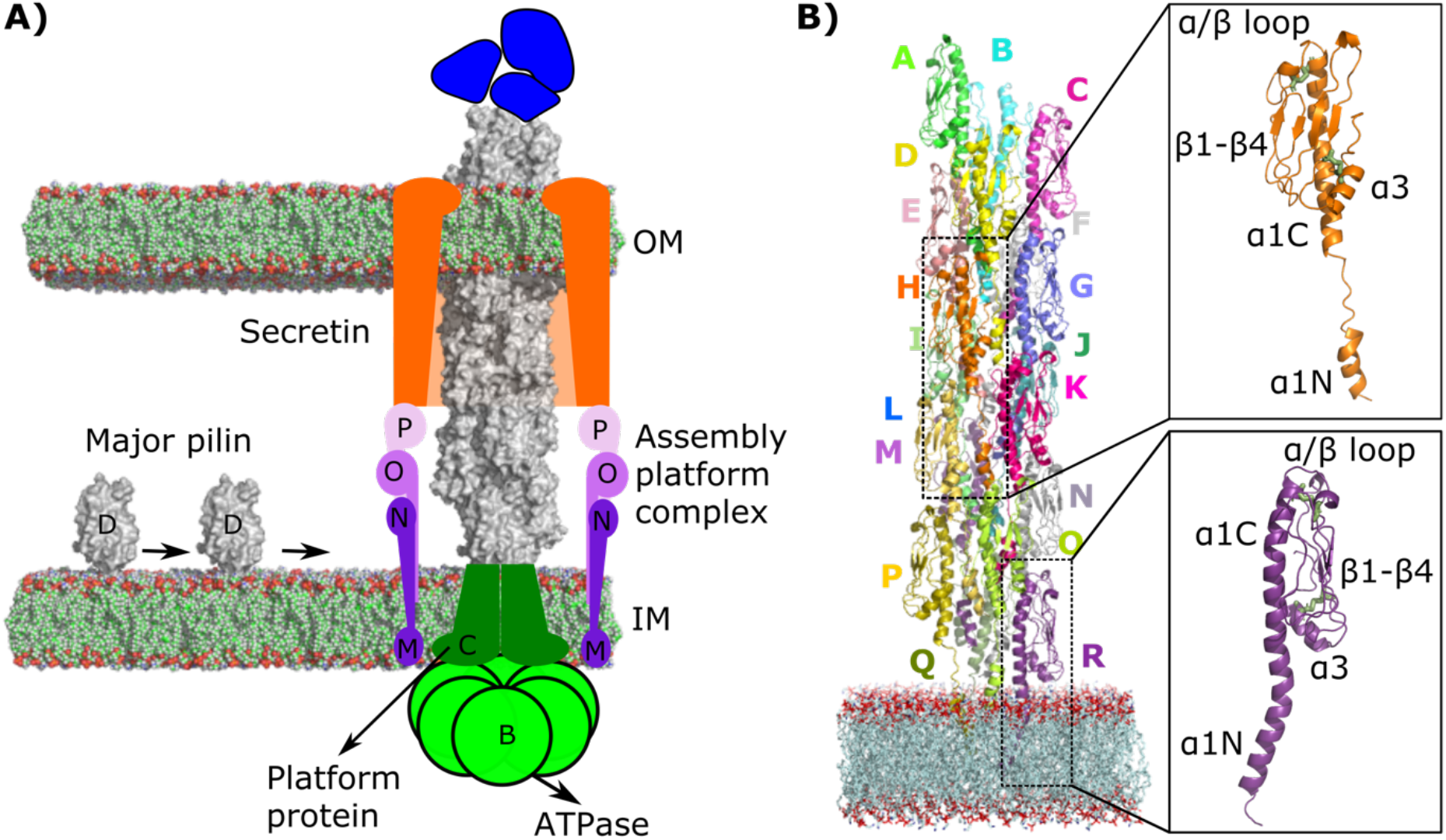
Schematic representation of the EHEC T4P assembly system. (A) Major pilin PpdD (gray) and minor pilins (blue) are depicted. Cytoplasmic ATPases (B, light green) transmit the conformational changes to the IM platform protein (C, dark green) and to the assembly complex (M, N, O, P, purple shades), which is connected with the secretin (orange) in the OM. (B) The starting structure for the MD simulations is depicted. T4P subunits (A-R) are shown as cartoon models in different colors. The POPE membrane atoms are shown with spheres colored in green (carbon), red (oxygen), and blue (nitrogen). The insets highlight the structure of two subunits with two conformations for the linker: as a coil (in orange), and modelled as a helix (in purple). The disulfide bonds are highlighted with green sticks.

T4Ps are present in Enterohemorrhagic *Escherichia coli* (EHEC), an important human pathogen. The major pilin PpdD (or HcpA) is one of the EHEC virulence factors (Xicohtencatl-Cortes et al. 2007) and its sequence is highly conserved in Enterobacteriaceae (Luna Rico et al. 2019). The structure of the periplasmic domain of EHEC PpdD determined by NMR (Amorim et al. 2014; Bardiaux et al. 2019), combined with the cryo-EM density map of the EHEC pilus at 8 Å resolution, led to an atomistic model of the T4P filament (Bardiaux et al. 2019). A variety of structural data is available for other T4P, with major contributions coming from cryo-EM (Wang et al. 2017; Kolappan et al. 2016; Hartung et al. 2011; Craig et al. 2006; Craig et al. 2003; Parge et al. 1995). Despite the recent progress in cryo-EM and integrative structural biology, the high flexibility of this family of fibers often limits the resolution of the structure. Modeling is therefore a necessary step in structure determination, even for the highest resolution cryo-EM maps obtained to date, such as those of the *Thermus thermophilus* T4P (Neuhaus et al. 2020). Moreover, given the T4P flexibility and ability to rapidly extend and retract, a single structural model is not sufficient to fully understand their behavior, nor to design new ways to regulate and interfere with their functions. A crucial ingredient currently missing is a comprehensive and accurate characterization of the dynamic properties of these systems.

The main objective of the present study is to investigate internal dynamics of T4P to be able to propose strategies that regulate T4P behavior by directly targeting the pilus. With the aim to reach a deeper understanding of T4P dynamics and its functional role, we performed a total of 108 µs all-atom molecular dynamics (MD) simulations of EHEC pili. This allowed us to reveal, at atomistic detail, the network of dynamic interactions between subunits and the role of different regions in modulating filament flexibility. This in-depth characterization of the conformational dynamics can also help explain the physical-chemical basis of the molecular mechanisms behind the filament formation. We also analyzed the effect of specific structural features of pilin subunits on pilus dynamics and stability. While most T4a pilins have one disulfide bridge, defining the C-terminal loop (Amorim et al. 2014), and homologous type II secretion pseudopilins have none, enterobacterial T4P has an additional, conserved disulfide bond connecting the N-terminal α helix with the globular domain (Luna Rico et al. 2019). In this study, by combining *in silico* analysis with mutagenesis and biochemical assays, we analyzed the effects of these disulfide bonds on pilin stability and dynamics, as well as on the assembly of the pilus. Finally, we studied the role of ions. Other filaments of the T4P superfamily, such as archaeal flagella and the type II secretion pseudopili, are stabilized by calcium or other cations (Meshcheryakov et al. 2019; Lopez-Castilla et al. 2017; Korotkov et al. 2009), however their effect on T4aP has not been investigated. Here, by using NMR we showed that PpdD subunit binds calcium and identified the residues involved in this interaction. We combined MD analysis with targeted mutagenesis and functional assays to characterize the interaction of calcium and other ions within the assembled pilus and understand their role in fiber biogenesis and dynamics. Together, our MD analysis and *ex vivo* and *in vitro* assays allowed us to identify the key structural elements required for the stability and assembly of T4P.

## RESULTS

We performed a series of all-atom MD simulations listed in **Table 1**. To elucidate the role of ions in pilus dynamics, we carried out MD simulations of T4P in the absence of ions and in the presence of Ca^2+^, Na^+^, Mg^2+^, Mn^2+^ and at different salt concentrations. The effect of Ca^2+^ was studied at different temperatures: 36.85°C (310 K), 60°C (333.15 K) and 80°C (353.15 K). We also evaluated the role of specific subunit features in pilus stability and function, notably the two disulfide bridges present in PpdD (by analyzing double substituted PpdD variants C50C60S and C118C130S), and the fully conserved residue E5 (variant E5A). Finally, we studied the effects of different histidine protonation states for the essential residue H54: *i)* HSE (ε), *ii)* HSD (*δ*), and *iii)* HSP (ε*−δ*). In total, we performed 108 *μ*s MD simulations of 16 different systems, with three replicates for each. For all analyses, we considered the union of the results from the replicates. A protonated ε nitrogen of H54, and a salt concentration of 100mM NaCl were assumed for all studied systems, unless stated otherwise. In the simulated pilus, the subunits were labeled from A (the most distant from the membrane) to R (the closest to the membrane) (**Figure 1B**). To reduce the edge effects, we excluded from the analyses the 4 top (A-D) and 4 bottom (O-R) subunits and only considered the 10 intermediate protomers (E-N), hereafter designated as “bulk” subunits.

**Table 1.** Details of the MD simulations. All studied systems, as well as their corresponding temperature, number of replicates and simulation time are reported here.

| System | Name | Temperature (K) | # Replicates | Simulation length (ns) |
| --- | --- | --- | --- | --- |
| Wild type | WT | 310 | 3 | 3000 |
| | $WT_{\text{Ca}^{2+}}$ | 310 | 3 | 3000 |
| | $WT^{(\delta)*}_{\text{Ca}^{2+}}$ | 310 | 3 | 3000 |
| | $WT^{(\epsilon-\delta)}_{\text{Ca}^{2+}}$ | 310 | 3 | 3000 |
| | $WT^{\text{NaCl-150mM}\#}_{\text{Ca}^{2+}}$ | 310 | 3 | 1000 |
| | $WT_{\text{Na}}$ | 310 | 3 | 1000 |
| | $WT_{\text{Mg}}$ | 310 | 3 | 1000 |
| | $WT_{\text{Mn}}$ | 310 | 3 | 1000 |
| Temperatures | $WT^{60^\circ}$ | 333.15 | 3 | 3000 |
| | $WT^{60^\circ}_{\text{Ca}^{2+}}$ | 333.15 | 3 | 3000 |
| | $WT^{80^\circ}$ | 353.15 | 3 | 3000 |
| | $WT^{80^\circ}_{\text{Ca}^{2+}}$ | 353.15 | 3 | 3000 |
| Substitutions | $C50C60S_{\text{Ca}^{2+}}$ | 310 | 3 | 2000 |
| | $C118C130S_{\text{Ca}^{2+}}$ | 310 | 3 | 2000 |
|  | E5A | 310 | 3 | 2000 |
| | $E5A_{\text{Ca}^{2+}}$ | 310 | 3 | 2000 |
| Total |  | - | 48 | 108000 |

\*The protonation of $\epsilon$ nitrogen for H54.
#The salt concentration of 100mM NaCl were considered for all the studied systems, unless stated otherwise.

### Impact of pilin flexibility on length fluctuations

T4P are highly flexible fibers that undergo substantial length fluctuations (Biais et al. 2010) and show remarkable resistance to force (Biais et al. 2008). We first studied in detail the relationship between individual residue fluctuations of EHEC pilin subunits and the overall flexibility of the pilus bulk subunit in MD simulations of wild-type pilus in the presence of calcium (WT_Ca_^2+^).

#### Residue fluctuations

The PpdD fold includes a long N-terminal α helix and a β-sheet globular domain (**Figure 1B**). The middle of the N-terminal helix (residues G11 to P22) is unstructured in assembled pilins and is hereafter referred to as the linker. Similar to most T4a pilins, a disulfide bond stabilizes the C-terminal loop (C118 -C130) of PpdD, in addition to a disulfide bond between residues C50 and C60, conserved in the enterobacterial T4aP subclass (**Figure 1B**). The dynamics of every region, calculated as the root mean square fluctuations (RMSF) of every residue with respect to the average conformation during the final 2.5 *µ*s from the three MD replicates, is shown in **Figure 2A**. The maximum fluctuations (*RMSF*_*max*_) mapped on the structure (**Figure 2B**) revealed several highly flexible regions (*RMSF*_*max*_ *>* 4 Å): *(i)* G62-V81 (α/β loop and β1), *(ii)* E92-N95 (β2*/*β3 loop), *(iii)* W105-T112 (β3*/*β4 loop), and *(iv)* D137-N140, the C-terminal region.

**Figure 2.**
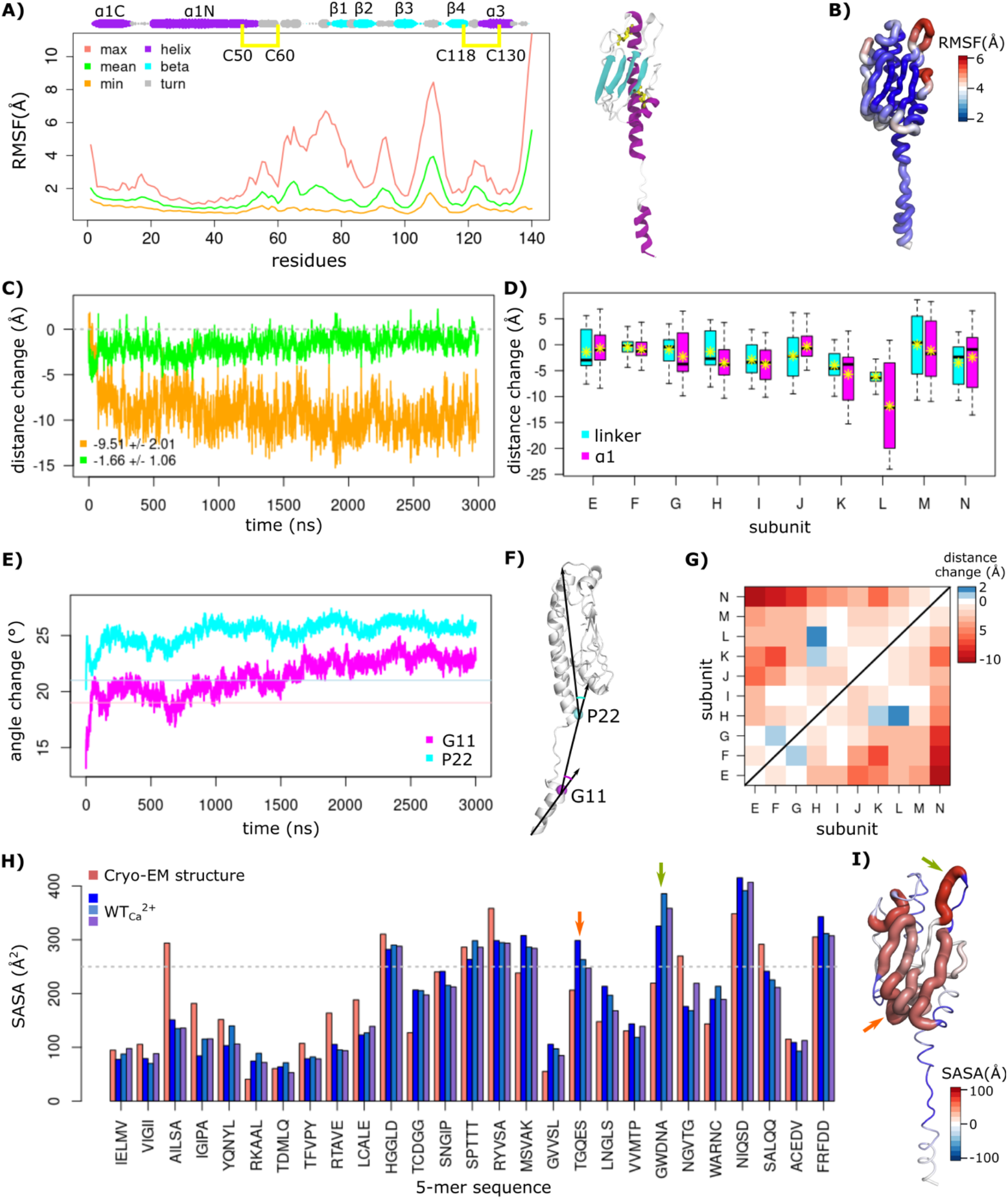
Residue fluctuations of the wild-type T4P from the MD simulations. (A) The per-residue RMSF of atomic coordinates are measured from the MD simulations, with respect to the average conformation. The maximum, minimum and average RMSF values were calculated over the bulk subunits from the three replicates and plotted as pink, orange, and green curves, respectively. The secondary structures are indicated (size of the rounds proportional to the persistence of the secondary structure along the MD trajectories). The starting structure of a single subunit is shown with the α helices and β strands colored in purple and cyan, respectively. The two disulfide bridges are shown as yellow sticks. (B) The maximum RMSF values are mapped on the structure. (C) The average filament distance change is measured over the three replicates for the full-length pilus (in orange), and bulk subunit (in green). (D) The changes of distances between the Cα atoms of the first and last residues of the linker (G11 and P22, in cyan), and α1 (F1 and E53, in magenta) are reported for bulk subunit with respect to their initial size, over the last 2.5 *μ*s of each replicate simulation. The average values for each subunit (letters E to N) are shown with yellow stars. (E) The changes of angles formed at the G11 (in magenta), and P22 (in cyan) are averaged over bulk subunit and the replicates. Their initial values from the cryo-EM structure are shown with horizontal lines. (F) The segments used for the calculations of angles in (E) are shown on the structure. (G) Changes of distance between the center of mass of bulk subunit globular domains (β1−β4) are averaged over the replicates. (H) The average SASA of 5-mers over the subunits of pilus are shown in shades of blue for the replicates, and pink for the initial structure (PDB code: 6GV9). The gray line corresponds to the threshold used for the predictions of exposed regions. The arrows point to the segments that become exposed. (I) The difference between average SASA from the MD and the initial structure is mapped on the structure, with colors from blue (buried) to red (exposed). Arrows indicate the loops ^89^TGQES^93^ (orange) and ^104^GWDNA^108^ (green) that are the most exposed to the solvent.

#### Filament dynamics

PpdD has the longest linker of any other pilus of this class, providing more flexibility to the filament (Bardiaux et al. 2019). To further explore its role, we investigated how the α1 helix and filament length changes correlate with the dynamics of the linker region. In principle, changes of filament length could be achieved by re-arrangements of the pilins as more or less rigid blocks, or by conformational changes within the pilins. We found that, while the filament length changes along the simulation were large, the changes of the bulk subunits were subtle, with an average length decrease of 1.7 Å (**Figure 2C**). We calculated all changes of distance as the difference between the instantaneous values along the MD simulations and the initial values in the cryo-EM structure. The changes of distance along the linker region and α1 (see **Methods**) in the three replicates along every subunit show an overall reduction of length for both the linker and α1 (**Figure 2D)**. We also measured the angles formed at the two ends of the linker region: G11 and P22 (see **Methods** for angle definition), and observed an overall increase along the simulations (**Figure 2E**), indicating bending of α1*N* and α1*C* (**Figure 2F**). Hence, the overall reduction of the filament length in the simulations is predominantly a consequence of the high flexibility of the linker and an increase of the angles formed between the two segments of α1 and the linker. The linker was the main region responsible for overall length fluctuations of the filament, with a correlation coefficient of 0.8 (**Figure S1)**. The overall reduction of length along the pilus was accompanied by a decrease in the separation between globular domains of the bulk subunit (**Figure 2G** and **Figure S1**). In T4aP, due to the helical symmetry, every subunit (S), makes contacts with six other subunits (S_*-*4_, S_*-*3_, S_*-*1_, S_+1_, S_+3_, S_+4_), along the right-handed 1-start, left-handed 3-start and right-handed 4-start helices (Bardiaux et al. 2019). We observed a larger reduction of the S distance to subunits S_+3_ and S_+4_ (**Figure 2G**). Consequently, the overall length reduction along the pilus long axis brings pilin subunits closer to each other, resulting in compaction and strengthening their contacts.

#### Exposed segments

To understand how pilus dynamics modifies solvent exposure of different regions during MD simulations, we divided the PpdD primary sequence into contiguous pentapeptide segments and compared their solvent accessibility (SASA, Å^2^) to the starting cryo-EM structure (**Figure 2H**). We classified segments with a SASA of greater than 250 Å^2^ as exposed. The results show that residues in the linker and α1 helix become more buried, while the residues of the globular domain β1*−*β4 become more exposed. The differences between the average SASA from MD simulations and the initial values from the cryo-EM structure were mapped on the pilin structure (**Figure 2I**). For two segments in particular –^89^TGQES^93^ and ^104^GWDNA^108^ – highlighted with red and green arrows, respectively, the average SASA increased strongly over the simulations, indicating that the β2/β3 and β3/β4 loops (**Figure 2I**) become more exposed to the solvent. Interestingly, the ^104^GWDNA^108^ peptide on the β3/β4 loop also showed the highest degree of fluctuations from the RMSF analysis (**Figure 2B**). Previous NMR studies identified the same region as highly dynamic in the PpdD monomer in solution (Bardiaux et al. 2019).

### Influence of Hydrogen-bonding network on filament formation

The analysis of MD trajectories in terms of inter-subunit Hydrogen-bonds (H-bonds) and salt bridges revealed the network of interactions between the subunits (**Figure 3)**. To analyze H-bonds, we recorded the inter-subunit interactions present for more than 50% of the simulation time in at least one replicate and that are observed between at least three pairs of subunits along the pilus symmetry (**Figure S2**). Salt bridges were studied by measuring distances between the center of mass of oxygen atoms from the acidic side chains and the center of mass of nitrogen atoms from the basic side chains, for all bulk subunits (see **Methods**). The results were mapped on one set of subunits within the helical symmetry (S_*-*4_ S_*-*3_ S S_+1_) (**Figure 4A**), where the thickness of the lines indicates the number of subunits for which the interactions are present. We observed 4 distinct pairs of H-bonds and salt bridges at the S-S_+1_ interface, 5 pairs at the S-S_+3_ interface, and 2 pairs at the S-S_+4_ interface. A strong network of H-bonds is present along the 1-start helix, involving mostly the α1 helix, whereas the majority of salt bridges are formed along the 3-start helix. Two sets of residues were pivotal for those interactions: *i)* K30, D35, E53, H54, R135, E92, E131, D132, involved in salt bridges, and *ii)* T2, L3, E5, L16, Y27, Q38, R74, N140, forming H-bonds. Some residues formed both salt bridges and H-bonds, *e*.*g*., D35-R135 and K30-E53. To experimentally validate the predicted importance of these interactions for pilus stability, we generated alanine substitutions and performed piliation assays. In our assay conditions, around 50% of total wild-type PpdD was assembled into pili on average (**Figure 4B**). In comparison, piliation was fully or partially impaired for a large number of these variants including T2A, E5A, L16A, Y27A, K30A, D35A, E53A, H54A, and R74A, supporting the role of these residues in assembly or stability of EHEC pili.

**Figure 3.**
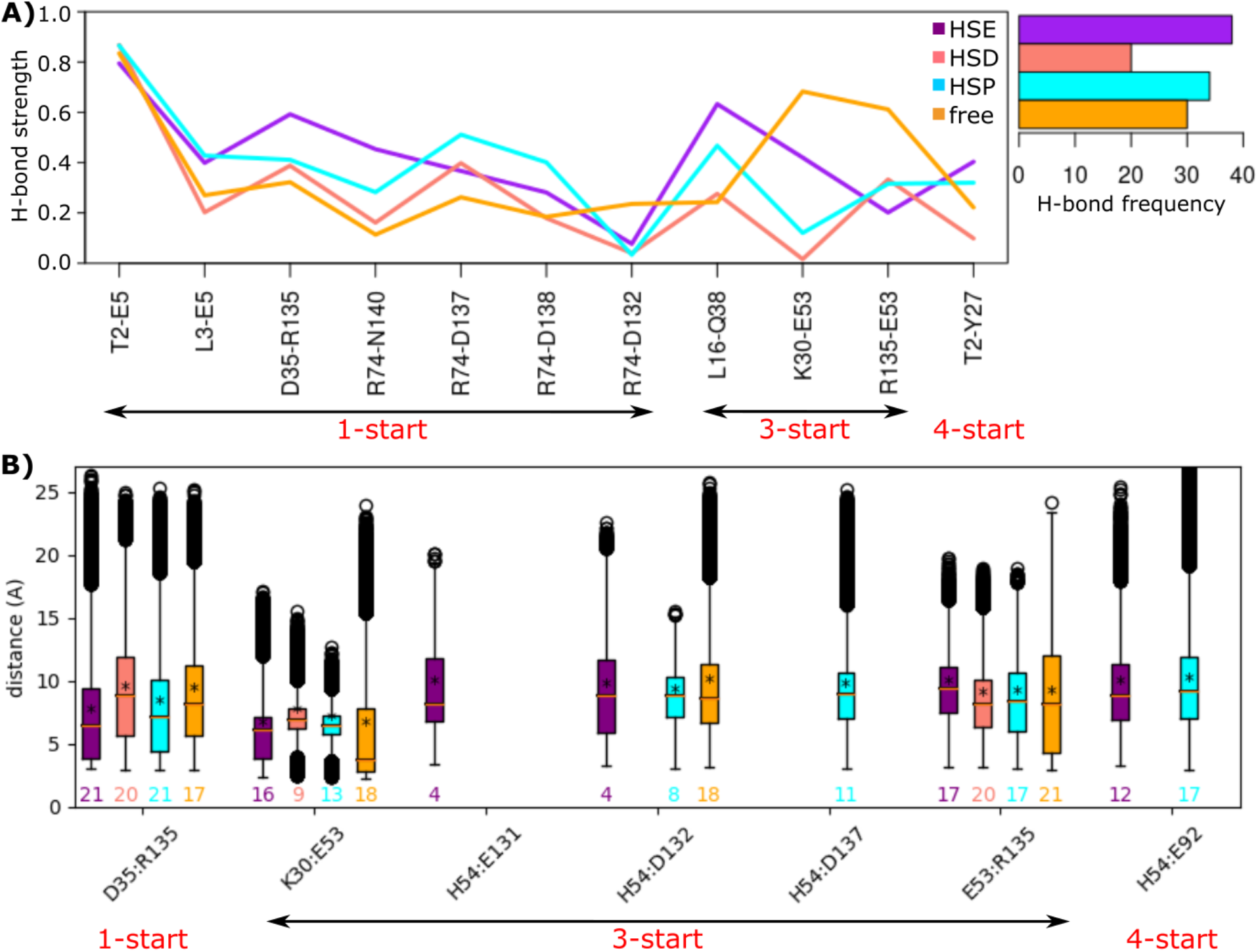
Inter-subunit interaction network. (A) The strength of H-bonds, computed as the percentage of conformations in which the H-bond is formed and the total number of H-Bonds formed along the pilus symmetry are reported for the calcium free and bound form pilus with different protonation states of His 54: HSE (ε), HSD (δ), and HSP (ε−δ). (B) The distances (Å) between pairs of residues forming salt bridges at the interface between subunits of the pilus are shown. The values above x-axis are the frequency of each salt bridge along the pilus symmetry. The left right arrows below each plot highlight the symmetry type of the interacting pairs. The data regarding the calcium bound (HSE, HSD, HSP) and free forms of the pilus are colored in orange, cyan, salmon and purple, respectively.

**Figure 4.**
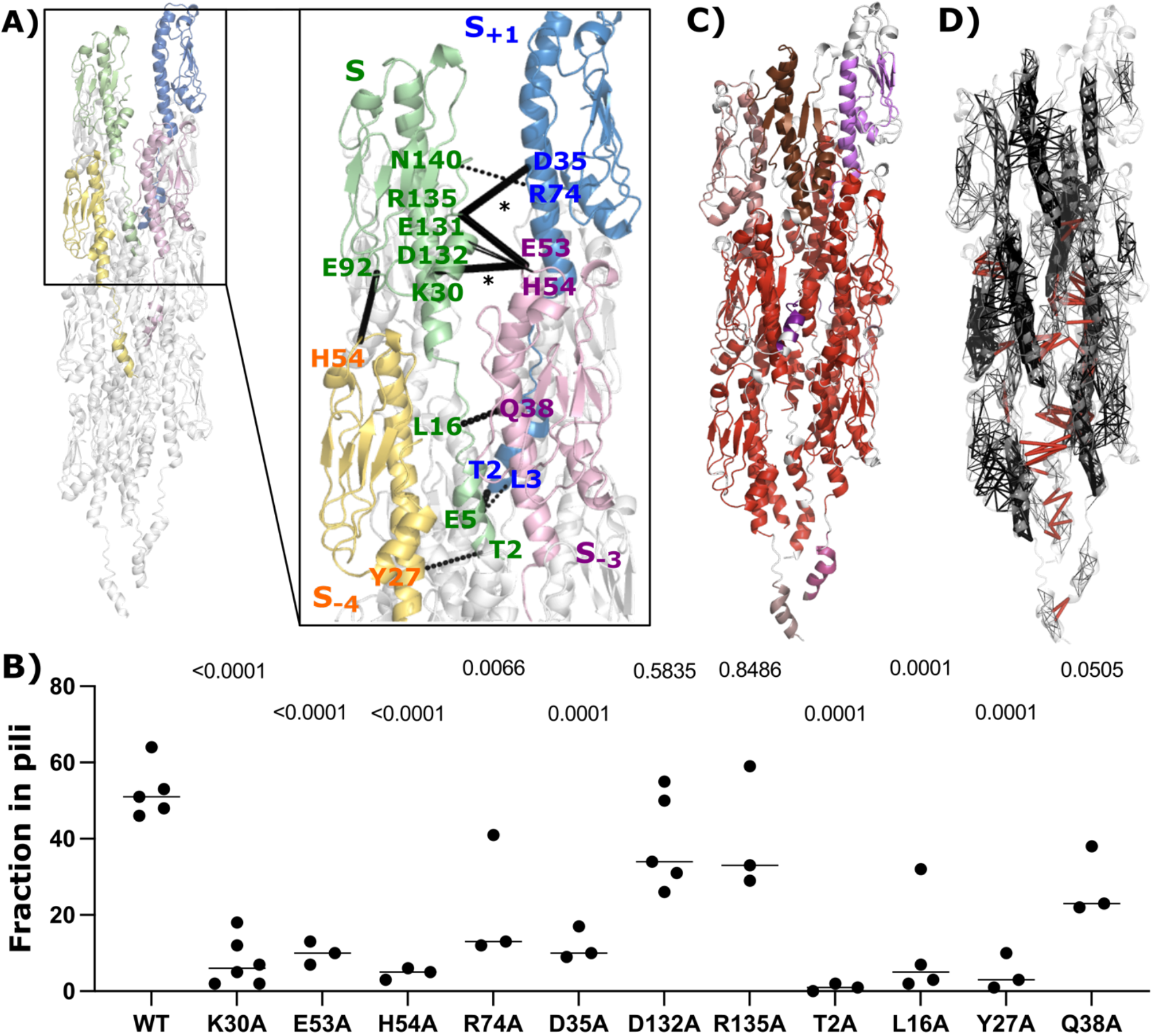
Inter-subunit interactions of the pilus. (A) Key residues involved in the interactions forming the pilus symmetry are highlighted on the first set of four protein subunits (S+1, S, S−3, S−4), colored in blue, green, pink and yellow, respectively. Dashed lines correspond to H-bonds and full lines to salt-bridges, where the thickness of the connections is proportional to their number of occurrences summed up according to the subunit symmetry along the pilus. (B) Dot plot comparing piliation efficiency of PpdD and its variants carrying alanine substitutions of individual residues. P values are shown above each graph as calculated in comparison with PpdD wild type piliation. (C) Pathways-based communication blocks (CBspath) identified by COMMA2 are mapped on the structure, with different colors. (D) The set of intra and inter-subunit pathways (> 3 residues) are displayed in black and red, respectively, as segments linking residues’ Cα atoms. The thickness of black segments is proportional to the number of pathways linking the residue pair.

We also measured the distances between all pairs of residues forming inter-subunit hydrophobic contacts for at least 70% of the simulation time (averaged over the helical symmetry) and recorded those within 3 Å (**Figure S3**). As expected, the majority of such residues belong to the α1 helix and form strong hydrophobic contacts.

Applying COMMA2 analysis (Karami et al. 2018) to the wild-type bulk subunit allowed us to identify a set of communication blocks *(CBs*^*path*^) in the pilus (see **Methods**). The residues within a *CB*^*path*^ are linked by communication pathways built by transitivity. A communication pathway is defined as a chain of residues displaying correlated motions and linked by stable non-covalent interactions, hence representing an efficient route to transmit information through physical interactions. The COMMA2 analysis revealed nine different *CBs*^*path*^ (**Figure 4C**), the largest of which (in red) contains residues from all 10 bulk subunits. To estimate the overall communication, we computed the number of pathways longer than 3 residues, and mapped intra-subunit (in black) and inter-subunit pathways (in red) on the structure of the pilus (**Figure 4D**). We found that the α1-helices play an important role in communications both within and between the subunits. Only 1% of the pathways constitute the inter-subunit communications (1547 pathways out of 202884), mainly involving the α1 helix residues. These results highlight the essential role of the α1 helix in pilus stability and allowed us to trace the communication route across the pilus.

### Calcium binding by PpdD modulates the stability of T4P

Calcium is important for function and stability of some filaments of the T4P superfamily, including type II secretion pseudopili and archaeal flagella (Meshcheryakov et al. 2019; Lopez-Castilla et al. 2017; Korotkov et al. 2009). So far, however, its effect on T4aP has not been investigated. Solution NMR analysis indicated weak calcium binding to the soluble periplasmic domain of the PpdD pilin (residues 26–140) as shown by chemical shift perturbations (CSP) in the presence of EGTA (a calcium chelating agent) or calcium (**Figure 5A**). Addition of EGTA resulted in the highest CSP values for residues L28, A31, ^51^ALEH^54^, F134 and F136. Among them, only E53 is negatively charged, located at the tip of the *α*1 helix, in a loop defined by the disulfide bridge C50-C60. As calcium was not added to any of our buffers, this indicates that it was present in the protein sample purified from the bacterial periplasm. Upon addition of up to 40 mM calcium, we observed the highest CSP for the same region, notably for residues A31, E53, H54 and F136. Based on these results, we had placed one calcium ion close to residue E53 in each PpdD subunit of the starting structure for the calcium-bound simulations (see **Methods**). Remarkably, during the simulations, we observed a movement of calcium toward the interface of three subunits: S, S_+3_, and S_+4_. We explored the interaction network in the vicinity of calcium, by comparing the calcium-bound and calcium-free simulations, and revealed a set of residues surrounding the calcium: E53, H54 (in the S protomer), K30, E131, D132, R135, D137 (in S_+3_), and D35, E92 (in S_+4_) (**Figure 5BC**). Interestingly, this set of residues overlaps with the set crucial for the interactions between subunits described above, forming H-bonds and salt bridges. Moreover, each calcium ion remained stable at the interface of three subunits and resulted in the formation of stronger H-bonds and salt bridges (**Figure 3**). In the calcium-free simulations, residue K30 (positively charged) interacts with residues E53 (negatively charged), through both H-bonds and salt bridges (**Figure 5B)**. However, the presence of calcium rearranged the interaction network due to its positive charge. In the calcium-bound simulations we observed that calcium oscillates between E53 (negatively charged) and D132 (negatively charged) as shown in **Figure 5C**. The distances between residues close to the calcium ion decreases overall by about 1 Å upon calcium binding, with an average distance of 7 Å for the calcium-bound simulation (HSE), compared to 8 Å for the calcium-free one (**Figure 5D**). While the distances decrease for most of the pairs, some exceptions are noted for K30-E53, R135-E53, and D132-H54. As observed for the salt-bridges and H-bonds, this can be explained by the fact that calcium intercalates between these residues. Although such rearrangements increase the distance between K30-E53 and R135-E53 and apparently weaken their contacts, these interactions are actually reinforced through the presence of Ca^2+^ ion that bridges those residues and neutralizes their charge. It is noteworthy that the side chain of H54 in its double protonation (HSP) state is closer to residue D137 (along the 3-start symmetry) and to E92 (along the 4-start symmetry), further strengthening the network of interactions. Importantly, functional assays show the importance of E53 and H54 residues for T4P assembly (**Figure 4B**).

**Figure 5.**
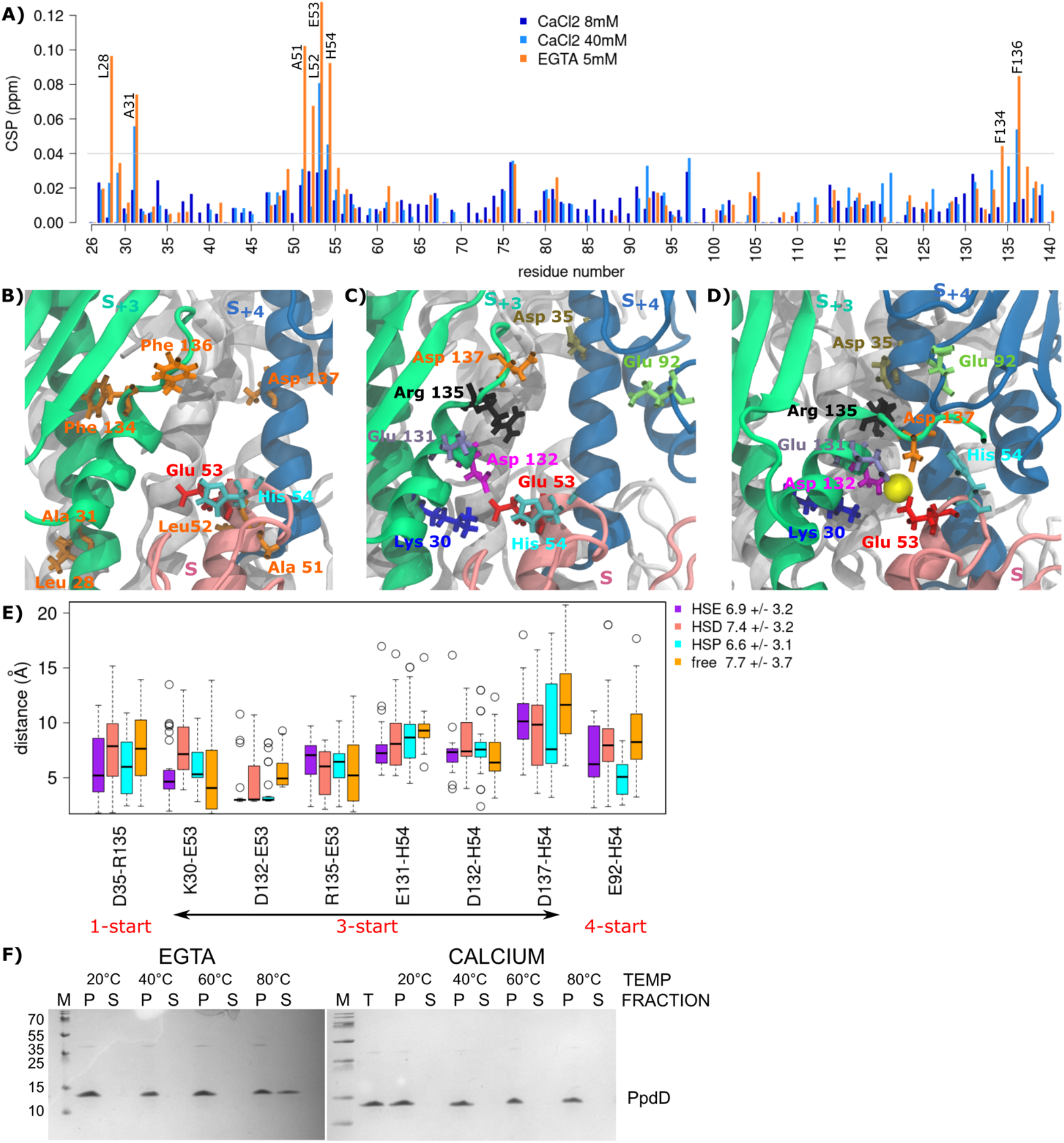
The effect of calcium binding on the overall interactions between subunits. (A) Histogram showing the CSP values of PpdD backbone amide signals in the presence of calcium (8mM in dark blue and 40mM in cyan) and EGTA (5mM in orange), as a function of residue number. Residues displaying highest CSP (> 0.04 ppm) are labeled. (B) Residues displaying significant spectral changes in the presence of EGTA are labeled on the structure and colored according to their physico-chemical properties (hydrophobic in orange, positively charged in blue and negatively charged in red). (C,D) Key residues involved in the interactions between subunits and calcium are highlighted as sticks in different colors. The three subunits (S, S+3, S+4), are colored in pink, green and blue, respectively, while the rest of the pilus are in white. The calcium ion is shown as a yellow sphere. (E) The comparison of distance variations between pairs of residues in the neighborhood of the calcium, in the absence of the calcium (free), and presence of calcium (HSE, HSD, HSP). (F) Thermal stability of the EHEC T4P. Aliquots of isolated pili were incubated for 5 min at 20°, 40°, 60° and 80°C in 50 mM HEPES 50 mM NaCl buffer supplemented with either 10 mM EGTA, or 20 mM calcium. Total pili samples (T) were fractionated by ultracentrifugation (see **Methods**). Equivalent volumes of pellet (P) and supernatant (S) fractions were analyzed by denaturing electrophoresis on 10% Tris-Tricin polyacrylamide gels and stained with Coomassie blue.

MD simulation analyses suggest that calcium should stabilize the pilus. To test this prediction, we designed biochemical experiments to assess the effects of calcium on pilus stability *in vitro*. Purified EHEC pili were incubated for 5 minutes at increasing temperatures in the presence or absence of calcium. Upon ultracentrifugation, intact pili were recovered in the pellet (P) and the dissociated subunits remained in the supernatant (S) (**Figure 5F**). The experiments showed that EHEC pili are quite resistant to temperatures up to 60°C, but at 80°C they start to disassemble *in vitro*. Pili incubated with buffer alone or buffer supplemented with EGTA behaved similarly, indicating a low level of divalent cation contamination in the buffer. Strikingly, the addition of calcium stabilized the pili at 80°C, fully preventing dissociation of PpdD from the filaments. To better characterize this effect, we carried out MD simulations at different temperatures (60°C and 80°C), in the absence and presence of calcium. While the pilus remained stable at both temperatures of 60°C and 80°C, these conditions induced high global fluctuations across the pilus (**Figure S4)**, and secondary structure perturbations specifically around the α3 helix (**Figure S5)**. Higher fluctuations were detected in the area of C50-C60 loop at 80°C compared to 60°C, possibly correlating with loss of fiber stability. However, for the simulations in the presence of calcium, we observed that the Ca^2+^ ions were not stable at their initial binding region and moved away after 50-200 ns in each simulation.

We also investigated the effects of ions and salt concentrations by performing MD simulations of T4P in 100 mM salt concentration by placing sodium (Na^+^), magnesium (Mg^2+^) or manganese (Mn^2+^) ions in the calcium binding site, or by increasing the salt concentration to 150 mM in the presence of calcium (Ca^2+^). In the simulations with 100 mM salt, the positively charged divalent ions behaved similarly to the calcium and remained stably bound in the same pocket, while in the simulations with 150 mM, the Ca^2+^ ions moved away from the binding site.

### Histidine 54 and the importance of its protonation

The single histidine (H54) at the tip of *α*1 helix, close to the calcium binding site is one of the key residues for the pilus formation, conserved in the enterobacterial T4aP subclass (Luna Rico et al. 2019). Its mutation to alanine abolished piliation (**Figure 4B**). To determine whether the protonation state of H54 affects pilus conformational stability, we performed three sets of MD simulations at different pH around the pKa of H54: *i)* HSE (ε), *ii)* HSD (*δ*), and *iii)* HSP (ε*−δ*). First, we measured the inter-subunit H-bonds over all the pairs of the bulk subunit, and recorded those that are present more than 50% of the simulation time in at least one replicate and between 3 pairs of subunits (**Figure 3A**). This analysis suggests that both HSE and HSP protonation states lead to stronger H-bond networks. However, the interactions are more persistent for HSE, occurring 38 times, compared to 34 for HSP. Interestingly, HSE results in the establishment of stronger H-bonds: (D35-R135) along the 1-start, (L16-Q38 and K30-E53) along the 3-start and (T2-Y27) along the 4-start symmetry. Also, the number and persistence of salt bridges are smaller for HSD (**Figure 3B**, the values above the x-axis). For both HSE and HSP, we observed salt bridges starting from H54 along the 3-start and 4-start symmetry. This analysis revealed a direct impact of H54 protonation state on stabilizing inter-subunit interactions, specifically along the 3-start and 4-start helical symmetry.

### The role of disulfide bridges and E5 on T4P structure and dynamics

#### Disulfide bridges

The presence of two disulfide bridges is conserved in enterobacterial T4aP subclass. Disrupting either of the two disulfide bridges led to a complete degradation of PpdD in bacteria, presumably due to increased exposure of dynamic loop regions to proteolysis in bacterial periplasm (**Figure 6A**). To assess the dependence of pilus stability on disulfide bonds, we treated purified pili with the reducing agent (DTT). Adding DTT had little effect on fiber stability at 60°C and below (**Figure 6B**). At 80°C, however, pili were fully dissociated, emphasizing the importance of these disulfide bonds. To gain molecular insight into their role, we performed MD simulations of pili lacking C50-C60 or C118-C130 disulfide bond. We observed that for both mutants, calcium ions were not stable and the fluctuations around the average increased with respect to the wild-type simulations, in particular close to the mutation sites (**Figure 6C**, the regions colored in red on the first two structures from the left). Similar perturbations were also observed at the level of secondary structures around the strands β1 for C50C60S_Ca_^2+^, and β4 for C118C130S_Ca_^2+^ (**Figure S5**). The mutations induced changes in interaction networks, with a strong decrease in the number of salt bridges, their persistency, and increase in the distances for the two mutants (**Figure 6D**). The only exception was the salt bridge between K30 and E53, which was more frequently present in the mutants compared to the wild type. This can be explained by the fact that the Ca^2+^ ions diffuse away in the simulations of mutants, allowing K30 and E53 to interact directly, compared to the wild-type simulations where this interaction was mediated by Ca^2+^ ion. Furthermore, the H-bond D35-R135 along the 1-start symmetry of the pilus became less stable, with the average strength of 0.59 for the wild type compared to 0.33 for the C50C60S_Ca_^2+^, and 0.28 for the C118C130S_Ca_^2+^ (**Figure 6E**).

**Figure 6.**
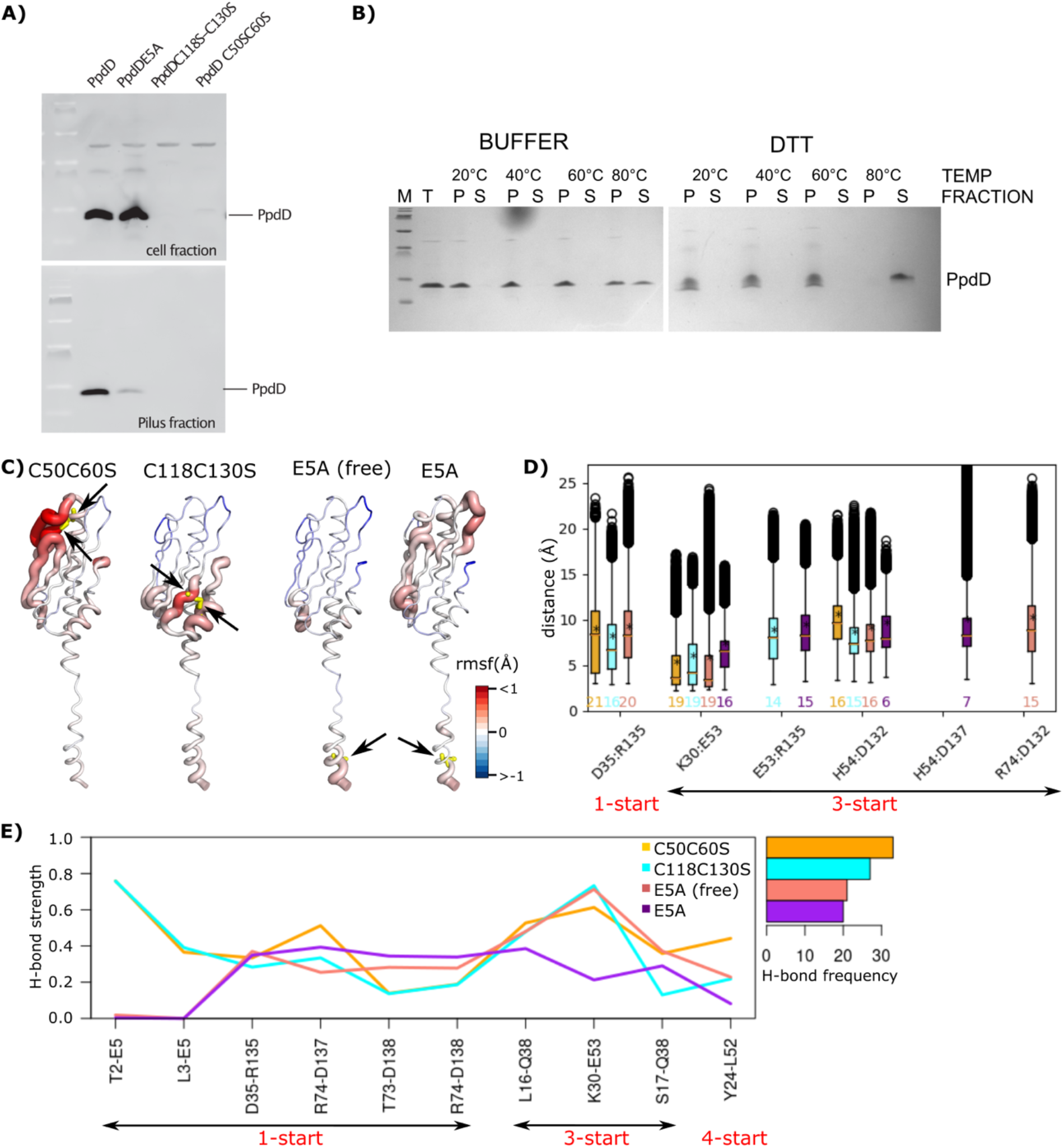
Role of disulfide bonds and E5 in pilus stability. (A) Piliation assay with single and double mutant variants. (B) Thermal stability of the EHEC T4P in the presence (BUFFER) and absence (DTT) of disulfide bridges. Total pili samples (T) were fractionated by ultracentrifugation and equivalent volumes of pellet (P) and supernatant (S) fractions were analyzed (see Methods). (C) The differences between the average per-residue RMSF are measured from the MD simulations of the C50C60S_Ca_^2+^, C118C130S_Ca_^2+^, E5A (free form), and E5A_Ca_^2+^ (from left to right), with respect to the wild-type (HSE) pilus. The differences are then mapped on the structure of each system. The mutation positions are shown with yellow sticks and pointed to by black arrows. (D) The distances between pairs of residues forming salt bridges, and (E) strength and frequency of H-bonds at the interface between subunits of the mutants are reported. The values above x-axis are the frequency of each salt bridge. The left right arrows bellow each plot, highlight the symmetry type of the interacting pairs.

#### Role of residue E5

The Alanine substitution of residue E5 prevents PpdD assembly (Bardiaux et al. 2019) (**Figure 6A**). We created the E5A variant *in silico* and carried out MD simulations in the absence and presence of calcium ions. For the simulations in the presence of calcium, we observed that Ca^2+^ ions were not stable, and moved away from their initial binding pocket. The comparison of local residue fluctuations with those of the wild-type pilus showed that the E5A substitution induces local perturbations around the fifth position (**Figure 6C**), as well as at the level of secondary structures in the *α*1N region (**Figure S5**). The mutation induced a major decrease in inter-subunit interaction strength and persistency (**Figure 6D**,**6E**). The total number of H-bonds along the pilus was reduced by about 50% (21 and 20 for E5A and E5A_*Ca*_, respectively, compared to 38 for wild type). On the other hand, intra-subunit interactions between E5 and F1 were increased from 51% in the simulations of the wild type to 82% and 74% in the E5A and E5A_*Ca*_ simulations, respectively. This can be explained by the disruption of the inter-subunit contacts of E5 along the 1-start helix.

## DISCUSSION

For this study of the conformational dynamics and stability of the EHEC T4P, we used a total of 108 *μ*s MD simulations of the wild-type T4P in different conditions, *i*.*e*., in presence and absence of different ions, protonation states, and salt concentrations, as well as point mutations, validated by experimental studies. It enabled us to identify regions responsible for the flexibility of the pilus, playing key roles for the overall length fluctuation of the filament. Moreover, our simulations revealed the network of interactions governing the communications across T4P, and allowed us to characterize a novel type of a calcium binding site that specifically stabilizes the assembled pilus polymer.

This study is to the best of our knowledge the most extensive study of the conformational dynamics and stability of a filament of this class. It is also the first study that uses a complete structural model with the “linker region” in the N-terminal helix. The loss of helical structure in α1N regions delimited by Gly and Pro residues was observed in the cryo-EM reconstructions of T4P from *N. meningitidis* (Kolappan et al. 2016), *N. gonorrhoeae* (Wang et al. 2017), *P. aeruginosa* (Wang et al. 2017), and EHEC (Bardiaux et al. 2019) in the 5-8 Å resolution range, in addition to two higher resolution structures for the wide and narrow forms of *T. thermophilus* with the resolutions of 3.2 Å and 3.5 Å, respectively (Neuhaus et al. 2020), and the structure of the homologous T2SS pseudopilus from *Klebsiella oxytoca* with a resolution of 5 Å (Lopez-Castilla et al., 2017). It has been proposed that this conserved feature of T4P facilitates the integration of pilin subunits into the pilus (Kolappan et al. 2016), allows T4P to withstand the extension under force, and to relax back to a native state when the force is removed (Wang et al. 2017). MD studies predating these structures, such as the atomistic and coarse-grain steered MD simulations to study the force-induced conformational changes of a T4P from *Neisseria gonorrhoae* (Zhao et al. 2017; Baker, Biais, and Tama 2013), were based on structural models of T4P with a continuous N-terminal helix. It is interesting that in those steered MD simulations, the pilus length variation was also linked to the particular region between G14 and P22 in the subunit. Among all the known structures of pili, the EHEC T4P has the longest inker region in the α1 helix, extending from residues G11 to P22. Our analysis identified in particular the role of the linker region in fiber flexibility: changes of its length show a strong correlation (0.8) with the overall flexibility of the filament. Such high correlation could explain the role of the linker in the filament resistance under force.

Despite the fact that in our simulations the pilus was in the “resting state” and absence of any forces, we observe an overall reduction of length at different levels, in the linker, α1 helix, globular domains and consequently along the pilus. Residues in the linker and α1 helix become more buried, while the residues of the globular domain β1*−*β4 become more exposed. This is consistent with the decrease of separation between the subunits, resulting in residues along the α1 helix becoming more buried. The reason for the length reduction in our simulations could be that we only simulate a small fraction of the pilus. Forces are typically generated by adding or removing pilin subunits, respectively, at the base of the pilus, leading to the extension or retraction of the pilus. These extension-attachment-retraction cycles are powered by the strongest linear motors (ATPase) known to date, and T4Ps have remarkable resistance to forces in the range of 100 pN (Clausen et al. 2009; Maier et al. 2002; Merz, So, and Sheetz 2000). It was also shown that the reversible force-induced conformational changes revealed the exposure of a hidden epitope (Biais et al. 2010), and the resistance to force was linked to the packing of α1 helices, maintaining their hydrophobic contacts within the core of the pilus (Baker, Biais, and Tama 2013). The N-terminal half of the α1 helix is highly conserved in the T4P, and is almost completely made up of hydrophobic residues, with the exception of T2 and E5. Our simulations highlight the role of hydrophobic contacts between the α1-helices in holding the core of the pilus together. However, they also underline the essential role of polar residues forming H-bonds (T2, L3, E5, L16, Y27, Q38, R74, N140) and salt bridges (K30, D35, E53, H54, R135, E92, E131, D132) at the interface of pilin subunits, along the 1-, 3-, and 4-start helix. Our results are in agreement with previous charge inversion mutagenesis analysis, which showed that the majority of charged EHEC T4P residues are required for pilus assembly (Bardiaux et al. 2019). At the same time, normal mode analysis of EHEC T4P indicated the role of interactions along the 3-start helix in pilus rigidity (Bardiaux et al. 2019). Our study highlights the crucial interplay of the linker region in α1, whose intrinsic flexibility induces long range dynamics along the filament, while the hydrophobic and polar interactions between the tightly packed helices provide stability.

In bacterial cells disulfide bridges are essential for PpdD monomer stability and protease resistance precluding the detailed analysis of their molecular role. MD data, coupled with the *in vitro* biochemical assays of an increase in temperature provided a consistent picture of the role of the disulfide bridges in the stability of T4P.

Strictly conserved residue E5, is crucial for T4P assembly, since it contributes to integration of pilin subunits into the pilus (Bardiaux et al. 2019; Nivaskumar et al. 2014; Lory and Strom 1997). The mutation E5A prevents the assembly at multiple levels (Luna Rico et al. 2019), and probably pilin extraction from the membrane as suggested by studies of the effects of E5A on the T2SS pseudopilin PulG (Santos-Moreno et al. 2017). Our MD data supports an additional role of this residue at the level of inter-protomer interactions, mostly by stabilizing the interactions between T2 and L3 on one subunit and E5 on the adjacent subunit, in agreement with previous work on other T4P (Craig et al. 2006; Wang et al. 2017).

MD simulations revealed the structural role of calcium in a member of the T4aP family, for which no classical calcium binding site had been described. By combining NMR chemical shift perturbation analysis, biochemical assays and MD simulations we revealed the molecular mechanism of calcium-mediated fiber stabilization. Simulations show how calcium bound at the tip of the pilin monomer becomes coordinated in assembled pili by additional residues from subunits S_+3_ and S_+4_, acting as a molecular glue. The set of residues involved in the interactions with calcium (K30, D35, E53, H54, E92, E131, D132, R135, D137) largely overlaps with the set of residues involved the in the network of H-bonds and salt bridges. We observed that calcium remains stable at the interface between pilin subunits, and results in the formation of stronger salt bridges and H-bonds at the interface. We also showed that the residues in the proximity of the calcium-binding region became more solvent exposed during the simulations. Complementary functional assays confirmed a subset of the residues in the calcium-binding region as critical for T4P assembly. In addition, *in vitro* biochemical assays showed that pili are resistant to temperatures up to 60°C and they start to disassemble only at 80°C. These results confirm previous findings regarding the thermal stability of meningococcal pili, also members of the T4aP subclass (Li, Egelman, and Craig 2012). Strikingly, the addition of calcium stabilized pili even at 80°C, fully preventing the dissociation of PpdD from the assembled filaments. Even though the simulations were too short to capture pilus dissociation, large fluctuations and secondary structure perturbations are in line with the biochemical data on fiber stability. Simulations with different ions support the requirement of positively charged divalent ions for stable binding.

The importance of calcium and the identification of a metal binding pocket at the interface between pilins in the simulations raises the possibility that this pocket could be exploited for drug discovery, since small molecules could bind there, affecting T4P assembly or function. T4P are virulence factors in many human pathogens. Newly emerging anti-virulence strategies have been developed to target the ATPases of T4P at the cytoplasmic base, resulting in disassembly of the pilus (Denis et al. 2019; Duménil 2019). Targeting the pilus directly rather than the basal body would have the advantage that potential drugs do not need to traverse the bacterial membrane.

This extensive characterization of the conformational landscape of T4P allowed us to reveal, at atomistic detail, the network of dynamic interactions between subunits and the role of different regions in modulating the filament flexibility. Such characterization of the conformational dynamics can also explain the physico-chemical basis of the molecular mechanisms behind the fluctuations in the overall length of the pilus. This valuable information has a profound impact at multiple different levels. From a fundamental point of view, our study elucidated the physico-chemical interactions that drive the behavior of large and dynamic molecular assemblies. These interactions must provide an optimal balance between structural integrity of the filaments and the flexibility required to perform their biological functions and to resist under stress. Furthermore, the acquired knowledge will guide rational strategies to modulate the behavior of these filaments that are critical for a variety of bacterial functions related to virulence.

## METHODS

### PPDD CALCIUM BINDING MONITORED BY NMR

The soluble periplasmic domain of EHEC PpdD comprising residues 26–140 of the mature protein, was produced and purified as previously reported (Bardiaux et al. 2019; Amorim et al. 2014). NMR data were acquired at 293 K on a Varian spectrometer operating at a proton frequency of 500 MHz. Proton chemical shifts were referenced to 2,2-dimethyl-2-silapentane-5 sulfonate (DSS) as 0 ppm. ^15^N were referenced indirectly to DSS (Wishart et al. 1995). The pulse sequences were from VnmrJ Biopack. NMR data were processed with NMRPipe/NMRDraw (Delaglio et al. 1995) and analyzed with the CcpNmr Analysis software package (Vranken et al. 2005). ^1^H–^15^N HSQC experiments were acquired on 0.4 mM PpdD in 50 mM Tris-HCl pH 7, 50 mM NaCl supplemented with either EGTA at 5mM or CaCl_2_ from 8-40 mM. Chemical shift perturbations (CSPs) of PpdDp backbone amide cross-peaks were quantified by using the equation CSP = [Δ *δ*H^2^ + (Δ *δ*N× 0.159)^2^]^1/2^, where Δ *δ*H and Δ *δ*N are the observed ^1^H and ^15^N chemical shift changes between the two experimental conditions. The residues belonging to the tag were not considered for this analysis. The ^1^H and ^15^N resonance assignments were from (Amorim et al. 2014).

### PILUS THERMAL STABILITY ASSAY

The *E. coli* strain PAP5386 is a *fimAB::kan ΔfliC* derivative of BW25113 F’*lacI*^*Q*^ (Datsenko and Wanner 2000). Bacteria were transformed with plasmids pMS41 encoding EHEC T4P assembly system and pCHAP8565 carrying the major pilin gene *ppdD* (Luna Rico et al. 2019). Bacteria were cultured for 5 days at 30°C on solid M9 glycerol agar plates containing 100 *μ*g.mL^-1^ of carbenicillin (Cb) and 25 *μ*g.mL^-1^ of chloramphenicol (Cm), supplemented with 1 mM isopropyl-β-D-1-thiogalactopyranoside (IPTG) to induce the expression of the T4P genes. Bacteria were harvested and pili were extracted by shearing and concentrated by ultracentrifugation as described previously (Luna-Rico, Thomassin, and Francetic 2018). To assess pilus stability, the equivalent amounts of pili were taken and resuspended in buffer (50 mM HEPES, 50 mM NaCl). The pilus samples were supplemented with 10 mM EGTA, 10 mM EGTA + 20 mM CaCl_2_, or 40 mM DTT, incubated at indicated temperatures for 5 min and then placed on ice. Intact pili were separated from dissociated subunits by a 30-min centrifugation at 53 000 rpm in the Beckman TLA55 rotor and table-top ultracentrifuge at 4°C. Equivalent amounts of total (T), pili-containing pellet (P) and supernatant fractions containing dissociated pilin subunits (S) were analyzed on 10% Tris-Tricin SDS-PAGE and stained with Coomassie blue.

### SITE-DIRECTED MUTAGENESIS

Mutations were introduced into the *PpdD* gene carried on plasmid pCHAP8565 by a modification of the QuickChange method for site-directed mutagenesis, as described previously (Bardiaux et al. 2019). The plasmids used in this study and mutagenic oligonucleotide primers are listed in **Tables S1** and **S2**.

### PILUS ASSEMBLY ASSAYS

To analyze pilus assembly, bacteria of *E. coli* strain BW25113 F’*lacI*^Q^ were transformed with plasmid pMS41 encoding the EHEC pilus assembly genes and pCHAP8565 or its mutant derivatives encoding PpdD variants (Luna Rico et al. 2019). Bacteria were cultured for 48-72 hours on M9 minimal agar plates (Miller 1972) containing Cb (100 *μ*g.mL-1) and Cm (25 *μ*g.mL-1). Bacteria were harvested and resuspended in LB medium at OD_600nm_ of 1. The suspension was vortexed vigorously for 1 min to detach pili from bacteria. Bacteria were pelleted by 5-min centrifugation at 16000 g at 4°C and resuspended in SDS sample buffer at a concentration of 10 OD_600nm_ per 1 mL. The supernatant containing pili was subjected to another round of centrifugation for 10 min. Pili were precipitated with 10% tri-chloroacetic acid on ice for 30 min and pelleted by a 30-min centrifugation at 16000g at 4°C. Pili pellets were washed twice with ice-cold acetone, air-dried and resuspended in SDS-sample buffer at a concentration equivalent to that of the bacteria. The equivalent volumes of each fraction were analyzed by denaturing SDS polyacrylamide gel electrophoresis (SDS-PAGE) on 10% polyacrylamide Tris-Tricine gels (Schägger and Von Jagow 1987). Proteins were transferred on nitrocellulose by Western blot and probed with anti-PpdD antibodies and secondary anti-rabbit antibodies coupled to HRP as described (Luna Rico et al. 2019). The fluorescent signal was recorded with a Typhoon FLA-9000 scanner (GE) and quantified by ImageJ. Data were processed and statistically analyzed with GraphPad Prism8 software.

### MOLECULAR DYNAMICS SIMULATIONS

#### Studied systems

The 3D coordinates of EHEC T4P were retrieved from the Protein Data Bank (Berman et al. 2000) with PDB code: 6gv9, residues 1 to 140, 8 Å resolution (Bardiaux et al. 2019). The published structure contains 14 subunits of the major pilin PpdD; each subunit comprises *(i)* a long N-terminal helix (α1) that is broken into two helices (α1*N* and α1*C*) by an extended linker from G11 to P22, *(ii)* four β strands forming a β sheet (β1*−*β2*−*β3*−*β4), *(iii)* a long loop (α/β loop) connecting the α1 to the β domain, and *(iv)* another helix (α3) at the C-terminus (**Figure 2A**). Two disulfide bonds are present in each subunit, between Cys residues (C50, C60) and (C118, C130). We performed the MD simulations of T4P in a 1-palmitoyl-2-oleoyl-sn-glycero-3-phosphoethanolamine (POPE) model membrane. The protein was initially placed near the membrane with the linker and N-terminus of the first subunit partially inserted into the bilayer (**Figure 1B**). The number of subunits was increased from 14 to 18 by applying the internal helical symmetry parameters. Moreover, the environment of the histidine (H54) was manually checked, and consequently protonated with a hydrogen at the ε nitrogen. We also performed MD simulations at two other protonation states: *(i)* a protonated δ nitrogen, and *(ii)* two protonated ε and δ nitrogens. In addition, a single substitution (E5A) and two double substitutions (C50C60S and C118C130S) were studied *in silico* using the CHARMM-GUI server (Jo et al. 2008).

#### Ion placement protocol

We identified the calcium binding site using NMR and by titration of T4P with calcium (see **PpdD calcium binding monitored by NMR**). From this experiment four consecutive residues, ^51^ALEH^54^ showed the highest chemical shift perturbation, among which only E53 is negatively charged. Consequently, we placed the calcium ion (Ca^2+^) close to this residue, and performed minimization. In addition to calcium, we evaluated the effect of other ions, namely: sodium (Na^+^), magnesium (Mg^2+^) and manganese (Mn^2+^). These ions were placed at the same position than the one chosen for Ca^2+^

#### Preparation

All systems were prepared with CHARMM-GUI membrane builder server (http://www.charmm-gui.org/?doc-1?4input/membrane) and the CHARMM36m force field parameter set (Huang et al. 2017; MacKerell Jr, Feig, and Brooks 2004): *(i)* hydrogen atoms were added, *(ii)* the ions were positioned at the tip of α1*C*, following the procedure mentioned in the previous paragraph, *(iii)* the POPE model membrane was used to build the inner bacterial membrane, *(iv)* the solute was hydrated with a rectangular box of explicit TIP3P water molecules with a buffering distance up to 14 Å, *(v)* Na^+^ and Cl^*-*^ counter-ions were added to reproduce physiological salt concentration (100 mM and 150 mM solution of sodium chloride). The following simulations were carried out: *(i)* T4P in the presence and absence of calcium with different protonation states: *WT, WT*_*ca*_ ^2+^, *WT*(*δ*) _*ca*_^2+^, *WT*(ε*−δ*) _*ca*_ ^2+^, *(ii)* T4P in the presence of different ions (calcium, sodium, magnesium, manganese) and salt concentrations (100 and 150 mM): *WT*^*NaCl*150*mM* 2+^, *WT*_*Na*_, *WT*_*Mg*_, *WT*_*Mn*_, *(iii)* T4P in the presence and absence of calcium at different temperatures (60°C and 80°C): *WT*^60°^, *WT* ^60°^_*ca*_ ^2+^, *WT*^80°^, *WT* ^80°^ _*ca*_^2+^, and *(iv)* a set of single and double mutants of T4P: *E5A* ^2+^, *C50C60S* ^2+^ and *C118C130S* ^2+^. A salt concentration of 100mM NaCl was used for all systems and the ε nitrogen of H54 was protonated, unless stated otherwise. For each system, three replicates of simulations were performed. Typically, these systems are composed of *∼*295,000 atoms, in a rectangular water box with dimensions of *∼*100 Å *×* 100 Å *×* 308 Å. The details of all studied systems are reported in **Table I**.

#### Production of the trajectories

The GROMACS 2019.4 package was used to carry out all simulations (Abraham et al. 2015). The energy minimization was performed by steepest descent algorithm for 10000 steps, to minimize any steric overlap between system components. This was followed by an equilibration simulation in an NPT ensemble at 310 K, allowing the lipid and solvent components to relax around the restrained protein. All the protein and lipid non-hydrogen atoms were harmonically restrained, with the constraints gradually reduced in 6 distinct steps with a total of 0.375 ns. The particle mesh Ewald algorithm (PME) (Essmann et al. 1995) was applied to calculate electrostatic forces, and the van der Waals interactions were smoothly switched off at 10-12 Å by a force-switching function (Steinbach and Brooks 1994). Production runs were performed in the NPT ensemble. The time step was set to 2.0 fs, the temperature was kept at 310 K (except for the simulations at 333.15 K and 353.15 K), temperature was kept constant using the Nosé-Hoover thermostat (Melchionna, Ciccotti, and Lee Holian 1993) and a constant pressure of 1 atm was maintained with the Parrinello-Rahman barostat (Parrinello and Rahman 1981). The SHAKE algorithm (Kräutler, Van Gunsteren, and Hünenberger 2001) was used to freeze bonds involving hydrogen atoms, allowing for an integration time step of 2.0 fs. The PME method (Darden, York, and Pedersen 1993) was employed to treat long-range electrostatics. Half-harmonic potentials were applied at the tip of the pilus (subunit A-D), in order to prevent the dissociation of the tip. PLUMED 2.6.0 (Tribello et al. 2014) and the PLUMED-ISDB (Bonomi and Camilloni 2017) module were used to add lower and upper walls on the distance between the *C*α atoms of four pairs of residues (*T*45_*A*_-*L*52_*B*_, *T*45_*B*_-*L*52_*C*_, *T*45_*C*_-*L*52_*D*_, *L*49_*D*_-*K*30_*A*_), with a force constant of 1000 *kcal/(mol Å)*. The choice of residues was made according to their fluctuations, and those with low deviations were selected. Coordinates of the system were written every 10ps. For every system, two or three replicates were performed, starting with different initial velocities as reported in **Table 1**.

#### Stability of the trajectories

Standard analyses of the MD trajectories were performed with the *gmx* module of GROMACS 2019.4. All analyses performed in this study were applied to the “bulk” subunit, *i*.*e*. remaining 10 intermediate subunits, after excluding 4 subunits from the top (A-D) and bottom (O-R) of the pilus (E-N). The root mean square deviations (RMSD) of backbone atoms (*Cα, C, N, O*) from the initial frame were recorded along each replicate (**Figure S6**). Based on the RMSD profiles, we performed the subsequent analysis over the subset of simulations where the systems are fully relaxed, *i*.*e*. considering the last 900 ns, 1500 ns and 2500 ns for the simulations of 1000 ns, 2000 ns and 3000 ns, respectively. The per-residue root mean square fluctuations (RMSF) were computed over the backbone atoms (*Cα, C, N, O*), with respect to the average conformation (**Figure S6**). The secondary structures were assigned with DSSP (Kabsch and Sander 1983) and averaged over the replicates (**Figure S5**). All studied systems remained stable along the MD trajectories.

#### Stability of the trajectories

Standard analyses of the MD trajectories were performed with the *gmx* module of GROMACS 2019.4. All analyses performed in this study were applied to the “bulk” subunit, *i*.*e*. remaining 10 intermediate subunits, after excluding 4 subunits from the top (A-D) and bottom (O-R) of the pilus (E-N). The root mean square deviations (RMSD) of backbone atoms (*Cα, C, N, O*) from the initial frame were recorded along each replicate (**Figure S6**). Based on the RMSD profiles, we performed the subsequent analysis over the subset of simulations where the systems are fully relaxed, *i*.*e*. considering the last 900 ns, 1500 ns and 2500 ns for the simulations of 1000 ns, 2000 ns and 3000 ns, respectively. The per-residue root mean square fluctuations (RMSF) were computed over the backbone atoms (*Cα, C, N, O*), with respect to the average conformation (**Figure S6**). The secondary structures were assigned with DSSP (Kabsch and Sander 1983) and averaged over the replicates (**Figure S5**). All studied systems remained stable along the MD trajectories.

#### Salt bridges

We used VMD to detect salt bridges if the distance between any of the oxygen atoms of acidic residues and the nitrogen atoms of basic residues are within the cut-off distance of 3.2 Å in at least one frame (Humphrey, Dalke, and Schulten 1996). Moreover, for every salt bridge we recorded the distance between the center of mass of the oxygen atoms from the acidic side chains and the center of mass of the nitrogen atoms from the basic side chains. Then we merged the distances from the three replicates of each system, and calculated the average distance of every pair.

#### Hydrogen bonds

We identified hydrogen-bonds (H-bonds) with the HBPLUS algorithm (McDonald and Thornton 1994). H-bonds are detected between donor (D) and acceptor (A) atoms that satisfy the following geometric criteria: (i) maximum distances of 3.9 Å for D-A and 2.5 Å for H-A, (ii) minimum value of 90° for D-H-A, H-AAA and D-A-AA angles, where AA is the acceptor antecedent. For a given H-bond between residues *i* and *j*, an interaction strength is computed as the percentage of conformations in which the H-Bond is formed between any atoms of the same pair of residues (*i* and *j*). We merged the results from different replicates and assigned the maximum strength to every pair.

#### Distances and lengths

The length of the filament was measured as the distance between the center of mass of *Cα* atoms of residues A51-E53 in the first four subunits and the center of mass of *Cα* atoms of residues F1-K30 in the last four subunits. The length of the linker was defined as the distance between *Cα* atoms of G11 and P22, and the length of the *α*1 helix as the distance between *Cα* atoms of F1 and E53. The separation between the globular domains were measured as the distance between the center of mass of *Cα* atoms from the residues forming the β sheet (β1*−*β4) on each subunit. The changes of distance were calculated as the difference between the observed values along the MD simulations and the initial values from the cryo-EM structure.

#### Angles

We measured the angles at the two ends of the linker, *i*.*e*. G11 and P22. For that, we considered three segments over the *α*1 helix: *(i)* residues F1-I10, *(ii)* I12-I21, and *(iii)* A23-E53, and measured the angles between them. The angle at G11 was defined based on the lines that best fit to F1-I10 and I12-I21 segments. Similarly, the angle at P22 was calculated between the lines that fit best to the I12-I21 and A23-E53 segments.

#### Residue burial

The average solvent accessible surface area (SASA, Å) was measured for residues in the wild-type pilus with the *gmx sasa* module of GROMACS for the initial cryo-EM structure and over the replicates of MD simulations. We identified segments of 5-residue long (5-mer) and calculated the SASA of every 5-mer as the sum of corresponding residual SASA values.

## COMMA ANALYSIS

COMMA2 (Karami et al. 2018) was applied to the replicates of MD simulations, and communication blocks were extracted. COMMA2 identifies pathway-based communication blocks (*CBs*^*path*^), *i*.*e*. groups of residues that move together, and are linked by non-covalent interactions, and clique-based communication blocks (*CBs*^*clique*^), *i*.*e*. groups of residues close in space that display high concerted atomic fluctuations. *Communication pathways* and *independent cliques* are used to construct a colored graph *PCN*(*N, E*) defined by nodes *N* that correspond to the residues of the protein and edges *E* that connect residues adjacent in a pathway or belonging to the same clique. COMMA2 extracts connected components from the graph by using depth-first search (DFS) to identify the protein *dynamical units*. These units are referred to as “communication blocks” (see (Karami, Laine, and Carbone 2016) for formal definitions and detailed descriptions). *Communication pathways* are chains of residues that are not adjacent in the sequence, form stable non-covalent interactions (hydrogen-bonds or hydrophobic contacts), and communicate efficiently. *Communication efficiency or propensity* is expressed as (Karami, Laine, and Carbone 2016):

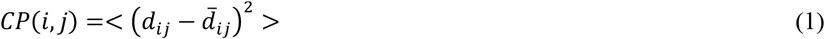

where *d*_*ij*_ is the distance between the C*α* atoms of residues *i* and *j* and 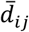 is the mean value computed over the set of conformations. Two residues *i* and *j* are considered to communicate efficiently if *CP*(*i,j*) is below a *communication propensity threshold*, CP_*cut*_. The strategy employed to set the value of *CP*_*cut*_ is explained in (Karami, Laine, and Carbone 2016). However, the algorithm is modified in this work by considering the definition of chains. Intuitively, it is expected that neighboring residues in the sequence forming well-defined secondary structure, communicate efficiently with each other. Therefore, we evaluate the proportion *p*_*ss*_ of residues that are in an *α*-helix, a β-sheet or a turn in more than half of the conformations. Then for every residue *i* surrounded by 8 sequence neighbors (4 before and 4 after), we compute a *modified communication propensity MCP*(*i*) as:

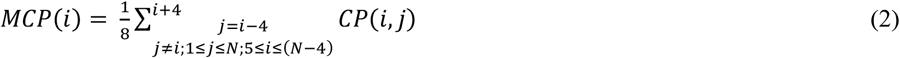

where *N* is the total number of residues in each chain. *CP*_*cut*_ is chosen such that the proportion *p*_*ss*_ of *MCP* values are lower than *CP*_*cut*_. Whenever more than one replicate of MD trajectories is available, we measured the *CP*_*cut*_ for each replicate and considered the average value for the identification of pathways.

## COMPUTATIONAL TOOLS

Trajectories generated by MD simulations were analyzed with gmx rms, gmx rmsf, gmx sasa utilities of GROMACS 2019.4 (Abraham et al. 2015). VMD (Humphrey, Dalke, and Schulten 1996) and PyMOL (DeLano 2002) were used for visualization and plots were generated using the R software package (Team 2013), and python (Van Rossum and Drake Jr 1995).

## LEAD CONTACT

Further information and requests for resources should be directed to and will be fulfilled by the Lead Contact, Michael Nilges.

## DATA AVAILABILITY

Raw MD trajectories and data are available from the authors upon request.

## ACKNOWLEDGMENTS

This work was funded by the Institut Pasteur, the Centre National de la Recherche Scientifique, the Fondation pour la Recherche Medicale (Equipe FRM 2017M.DEQ20170839114 to YK, ACL, NIP and MN), the French Agence Nationale de la Recherche (grant ANR-19-CE11-0020 to OF, NIP and MN), the Institut Pasteur Roux-Cantarini fellowship and NSERC grant (to JLT), Fulbright fellowship (to AO). YK would like to acknowledge the PRACE for awarding access to Piz Daint at CSCS, Switzerland.

## AUTHOR CONTRIBUTIONS

YK, NIP, OF and MN conceived the study. YK performed MD simulations and analyzed the data with BB, TM, NIP, OF and MN. ALC and NIP preformed NMR analysis. OF, JLT and AO analyzed pilus assembly and stability. YK, NIP, OF and MN wrote the paper. All authors read and approved the manuscript.

