## Supporting Materials for "Computational and biochemical analysis of type IV Pilus dynamics and stability"

### Supplementary Information.

Supplementary Tables 1.- 2.

Supplementary Figures 1.- 6.

**Supplementary Table 1. List of plasmids used in this study.**

| Plasmid name | Relevant characteristics | Source/reference |
|---|---|---|
| pCHAP8565 | p*lacZ-ppdD*, $Cm^R$, 15A *ori* | Luna-Rico et al, 2019 |
| pCHAP | placZ-ppdD E5A, $Cm^R$, 15A ori | Luna-Rico et al, 2019 |
| pMS1121 | placZ-ppdD T2A, $Cm^R$, 15A ori | This study |
| pMS1122 | placZ-ppdD L16A, $Cm^R$, 15A ori | This study |
| pMS1123 | placZ-ppdD Y27A, $Cm^R$, 15A ori | This study |
| pMS1079 | placZ-ppdD K30A, $Cm^R$, 15A ori | This study |
| pMS1053 | placZ-ppdD E35A, $Cm^R$, 15A ori | This study |
| pMS1124 | placZ-ppdD Q38A, $Cm^R$, 15A ori | This study |
| pCHAP6394 | placZ-ppdD E53A, $Cm^R$, 15A ori | This study |
| pCHAP6395 | placZ-ppdD H54A, $Cm^R$, 15A ori | This study |
| pMS1075 | placZ-ppdD R74A, $Cm^R$, 15A ori | This study |
| pMS1080 | placZ-ppdD D132A, $Cm^R$, 15A ori | This study |
| pMS1054 | placZ-ppdD R135A, $Cm^R$, 15A ori | This study |
| pCHAP6368 | placZ-ppdD C50S C60S, $Cm^R$, 15A ori | This study |
| pCHAP6367 | placZ-ppdD C118SC 130S, $Cm^R$, 15A ori | This study |

**Supplementary Table 2. Oligonucleotide primers used in this study.**

| Primer name/ mutation | Nucleotide sequence (5'-3') |
|---|---|
| PpdD T2A For | CAAGCAACGCGGTTTTgCACTTATCGAACTG |
| PpdD T2A Rev | CAGTTCGATAAGTGcAAAACCGCGTTGCTTG |
| PpdD L16A For | CATCATTGCCATTgcAAGCGCCATTGGTATTC |
| PpdD L16A Rev | GAATACCAATGGCGCTTgcAATGGCAATGATG |
| PpdD Y27A For | CGCTTATCAAAACgcCCTGCGCAAAGC |
| PpdD Y27A Rev | GCTTTGCGCAGGgcGTTTTGATAAGCG |
| PpdD K30A For | CAAAACTACCTGCGCgcAGCCGCACTCACCGAC |
| PpdD K30A Rev | GTCGGTGAGTGCGGCTgcGCGCAGGTAGTTTTG |
| PpdD D35A For | CAAAGCCGCACTCACCGcCATGCTACAAACCTTTG |
| PpdD D35A Rev | CAAAGGTTTGTAGCATGgCGGTGAGTGCGGCTTTG |
| PpdD Q38A For | CACCGACATGCTAgcAACCTTTGTGCCTTAC |
| PpdD Q38A Rev | GTAAGGCACAAAGGTTgcTAGCATGTCGGTG |
| PpdD E53A For | GAGTTGTGCGCGCTGGcgCATGGTGGATTAGATAC |
| PpdD E53A Rev | GTATCTAATCCACCATGcgCCAGCGCGCACAACTC |
| PpdD H54A For | GTTGTGCGCGCTGGAAgcgGGTGGATTAGATACC |
| PpdD H54A Rev | GGTATCTAATCCACCcgcTTCCAGCGCGCACAAC |
| PpdD R74A For | CCTACCACCACCgcCTATGTTTCAGCC |
| PpdD R74A Rev | GGCTGAAACATAGgcGGTGGTGGTAGG |
| PpdD D132A For | GCAAGCCTGCGAAGcTGTCTTCCGCTTTGATG |
| PpdD D132A Rev | CATCAAAGCGGAAGACAgCTTCGCAGGCTTGC |
| PpdS C50S For | CGTAGAGTTGTcCGCGCTGGAACATG |
| PpdD C50S Rev | CATGTTCCAGCGCGgACAACTCTACG |
| PpdD C60S For | GATTAGATACCTcCGACGGTGGCAGC |
| PpdD C60S Rev | GCTGCCACCGTCGgAGGTATCTAATC |
| PpdD C118S For | GGACGCGCAACTcCAATATTCAAAGTG |
| PpdD C118S Rev | CACTTTGAATATTGgAGTTGCGCGTCC |

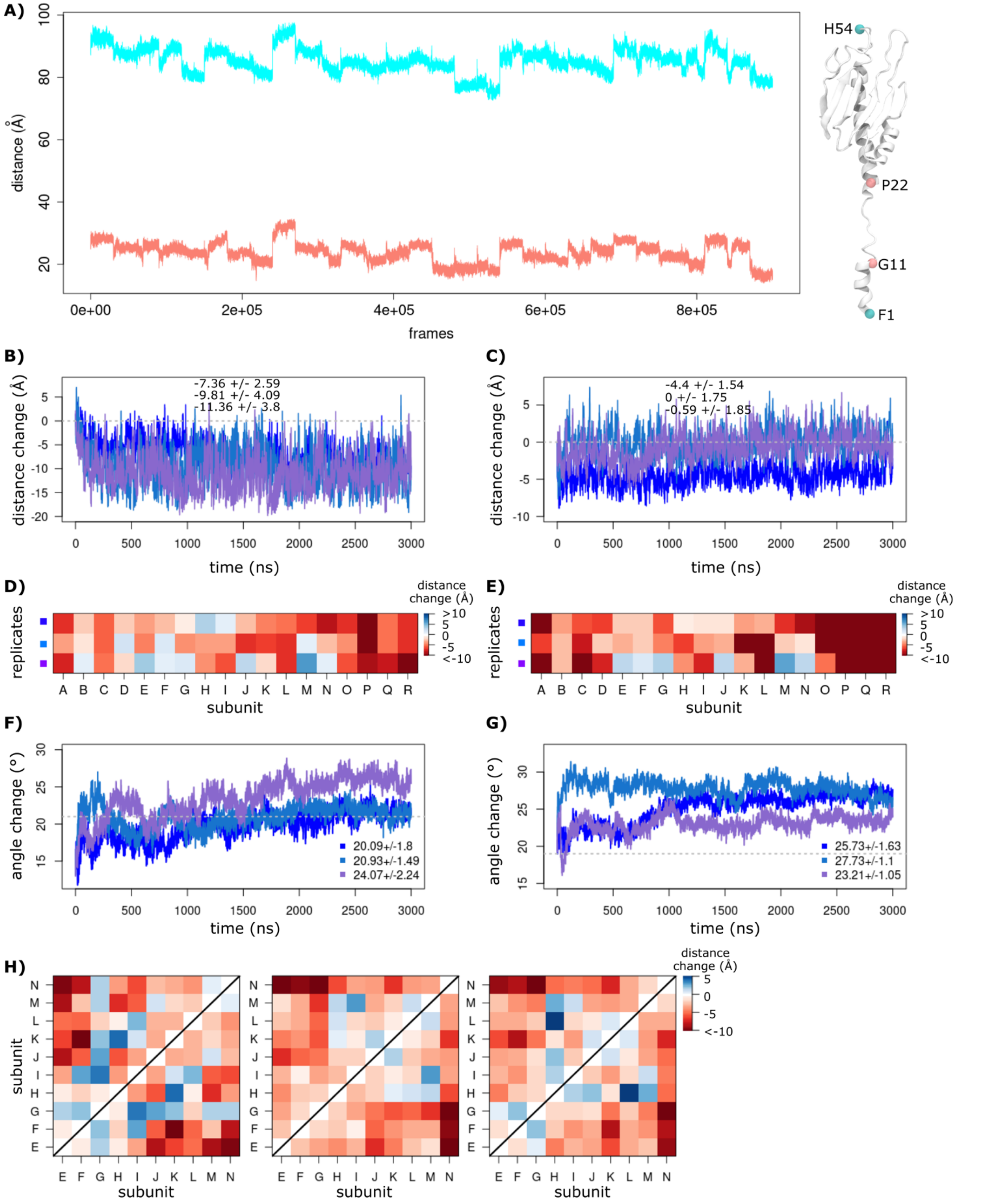


**Supplementary Figure 1. Distance changes along the simulation.** A) The distances between the two residues at both ends of the $\alpha 1$ helix (F1 and E53) and the linker (G11 and P22), were recorded along the simulations for all the subunits, and all replicates (shown in black and red, respectively). The average wild-type T4P filament distance change is measured along the MD simulations for the replicates of the B) full-length pilus, and C) bulk subunit. The changes of distances between the first and last residues of the D) linker (C$\alpha$ atoms of G11 and P22), and E) $\alpha 1$ (C$\alpha$ atoms of F1 and E53) are reported for each subunit with respect to their initial size, over the last 2.5 $\mu$s of each replicate. The changes of angles formed at the F) G11, and G) P22 are monitored for every replicate. Their initial values from the cryo-EM structure are shown with gray lines. H) The changes of distance between the center of mass of bulk subunit globular domains ($\beta 1-\beta 4$) is averaged for every replicate.

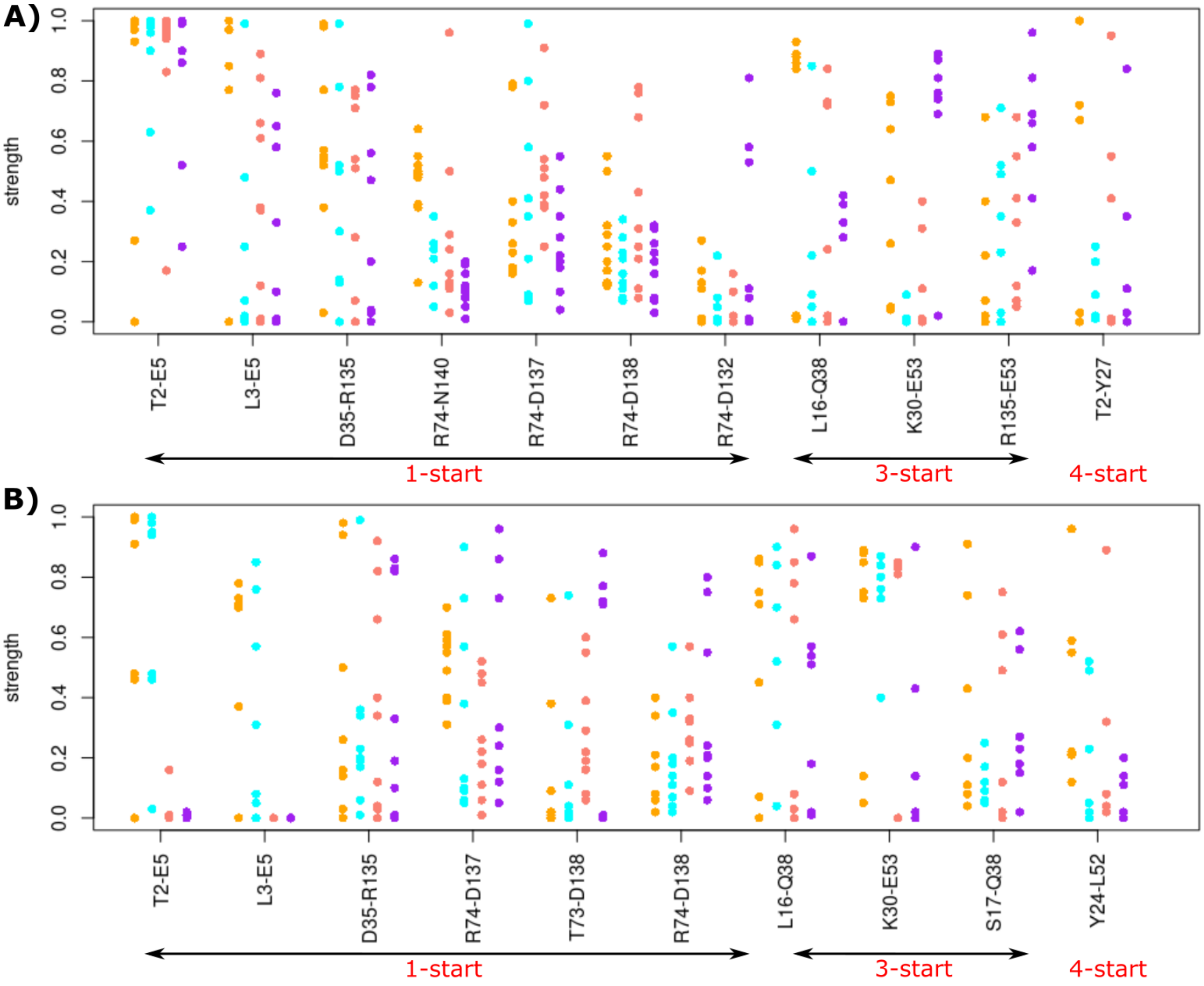


**Supplementary Figure 1. Details of the H-bonds from the MD simulations.** The persistency of all the H-bonds present for more than 50% of the simulation time is reported for the a) wild-type pilus (HSE, HSD, HSP and $Ca^{2+}$-free colored in orange, cyan, pink and purple, respectively), and b) mutants (C50C60S$_{Ca}^{2+}$, C118C130S$_{Ca}^{2+}$, E5A, E5A$_{Ca}^{2+}$, colored in orange, cyan, pink and purple, respectively). The values correspond to all pairs along the 3 replicates of bulk subunit for every studied system. The H-bonds are grouped according to the helical symmetry that they belong to.

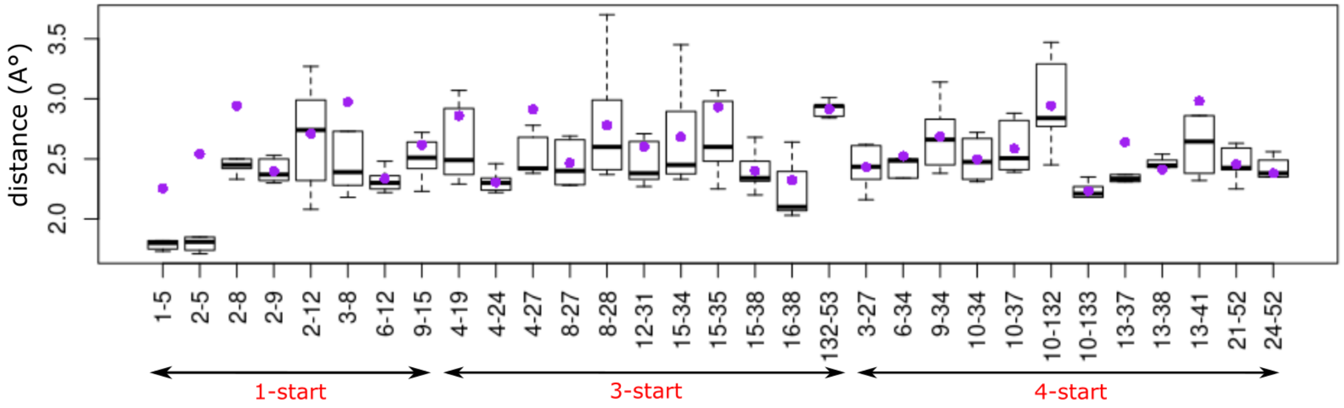


**Supplementary Figure 2. Inter-subunit interactions and distances of the pilus.** The changes of distances are recorded for all inter-subunit pairs of residues, showing an average distance of at most 3 Å and an average hydrophobic contact present at most 70% of the simulation time along the helical symmetry. The reported values correspond to the minimum distance and maximum interaction strength values over the replicates of the wild-type MD simulations. The purple circles are average distance values.

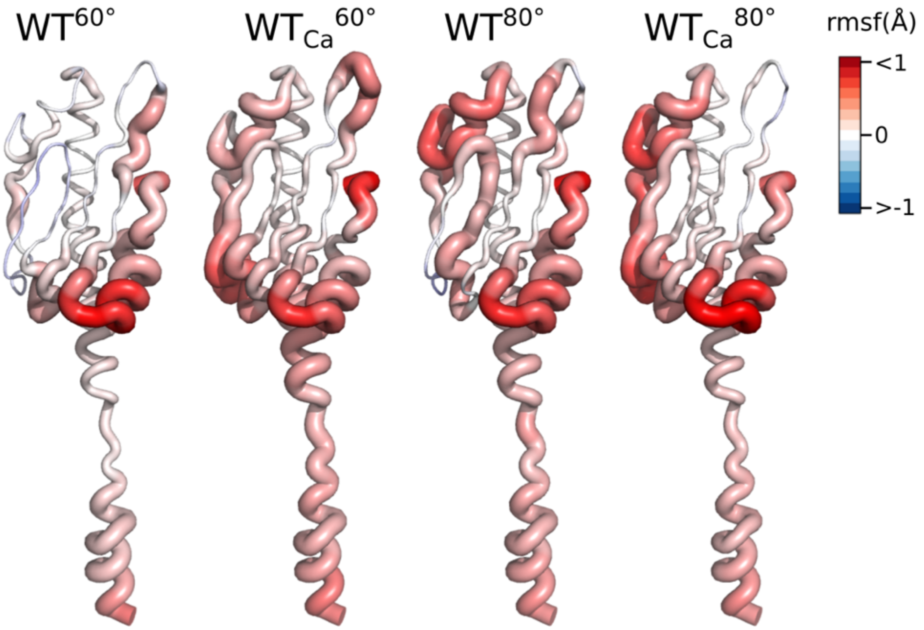


**Supplementary Figure 3. The effect of increase in temperature on the residue fluctuations.** The differences between the average per-residue RMSF are measured from the MD simulations of the $WT^{60ºC}$, $WT_{Ca}^{60ºC}$, $WT^{80ºC}$, and $WT_{Ca}^{80ºC}$ (from left to right), with respect to the wild-type (HSE) pilus. The differences are then mapped on the structure of each system. The color code represents the changes of fluctuations with the highest increase in red and decrease in blue.

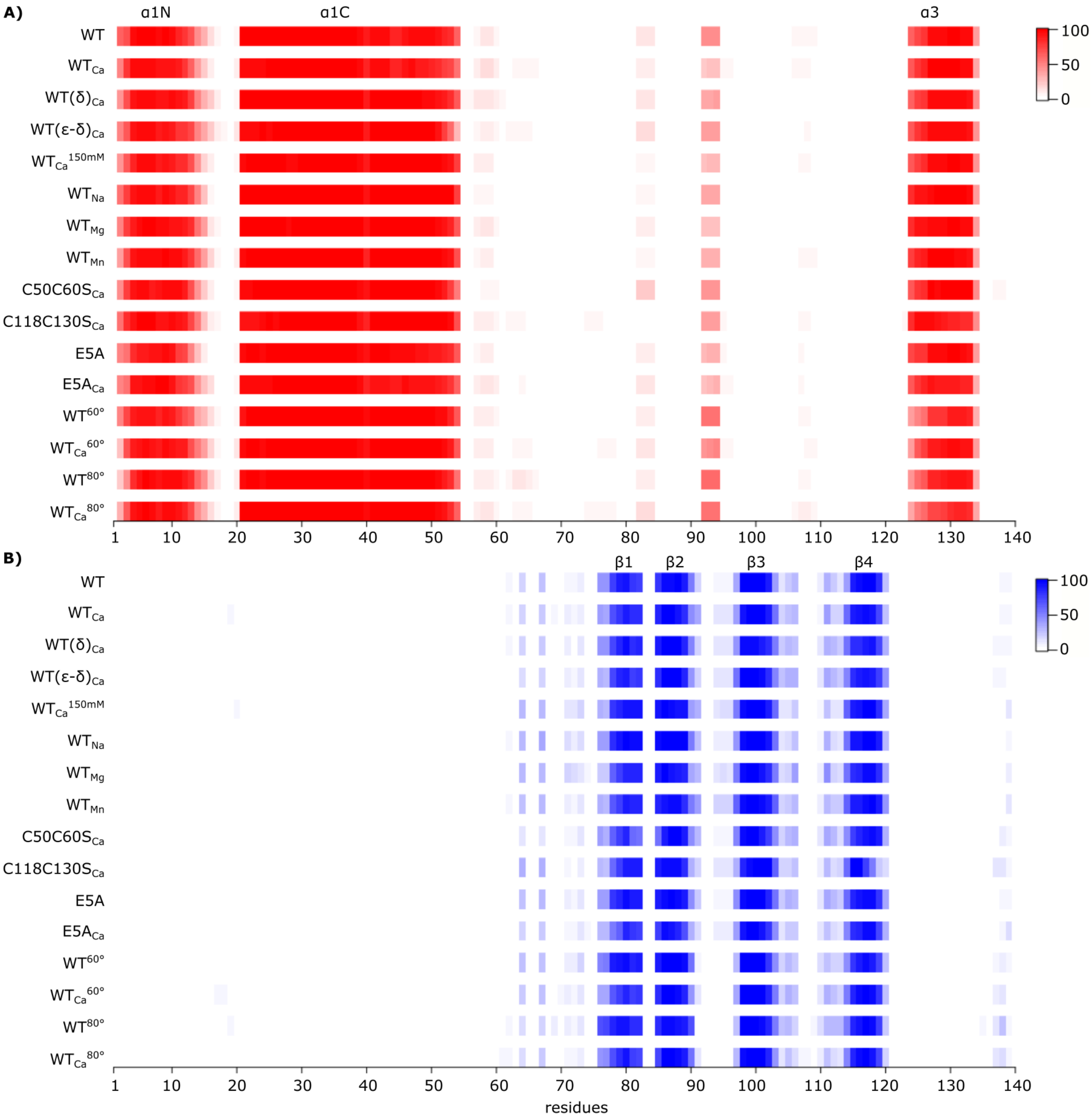


**Supplementary Figure 4. Secondary structure profiles for each studied system.** The stability of A) α helices, and B) β strands are recorded along each simulation and the average values are measured for every residue, over the replicates.

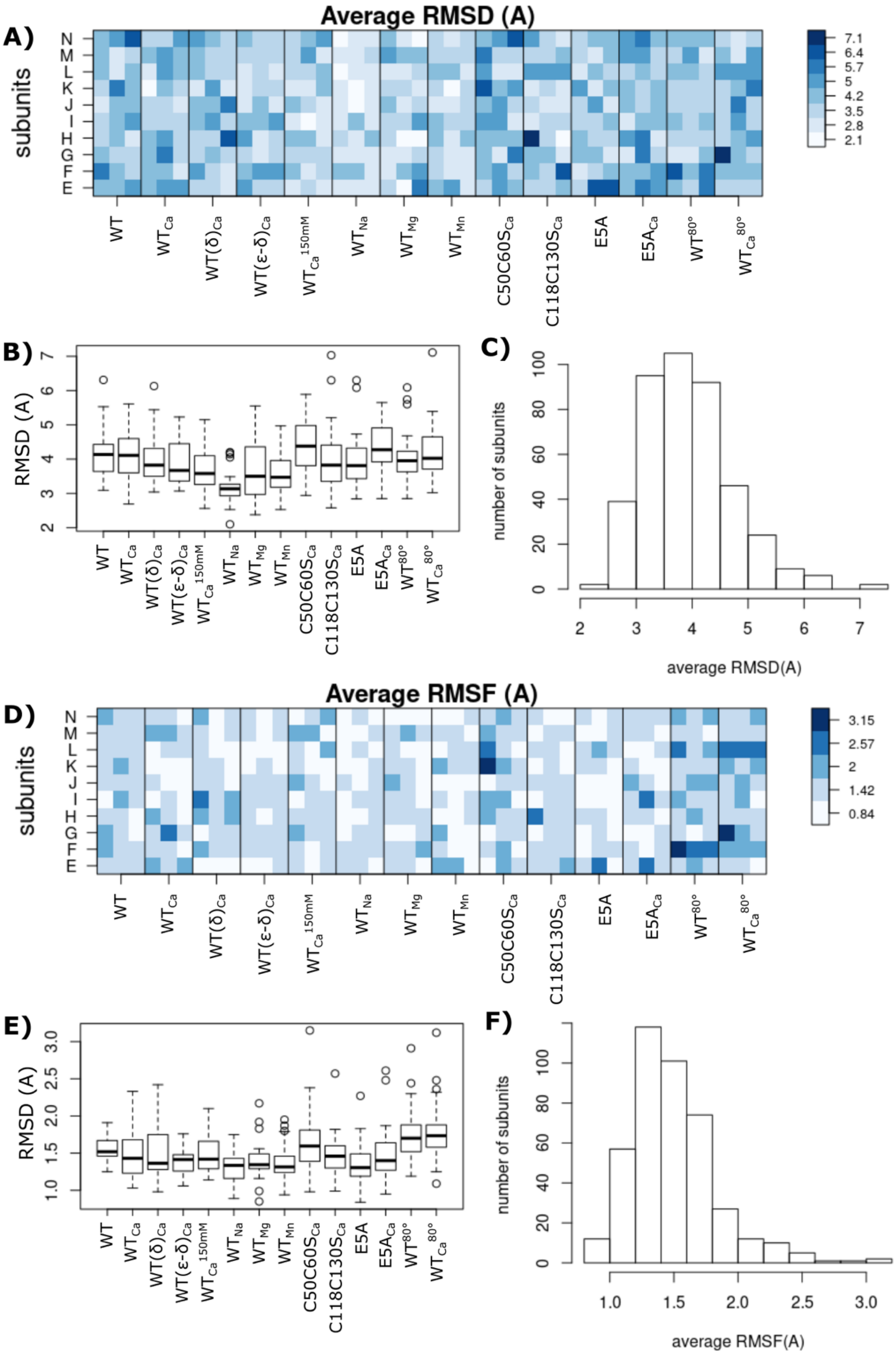


**Supplementary Figure 5. The average RMSD and RMSF of the bulk subunit for all the studied systems.** The RMSD of backbone atoms (Cα, C, N, O) from the initial frame were recorded along each replicate. A) The average RMSD value of each subunit (E-N) is reported for each replicate of every system. B) Every box corresponds to the average RMSD values of all subunits (from all replicates) of each system. C) The distribution of all the average RMSD values measured over each subunit of every replicate. The by-residue RMSF of backbone atoms (Cα, C, N, O) are recorded with respect to the average conformation for each replicate. D) The average RMSF value of each subunit (E-N) is reported for every replicate of every system. E) Every box corresponds to the average RMSF values of all subunits (from all replicates) of each system. F) The distribution of all the average RMSF values measured over each subunit of every replicate.
